# Meta-matching: a simple framework to translate phenotypic predictive models from big to small data

**DOI:** 10.1101/2020.08.10.245373

**Authors:** Tong He, Lijun An, Jiashi Feng, Danilo Bzdok, Avram J Holmes, Simon B. Eickhoff, B.T. Thomas Yeo

## Abstract

There is significant interest in using brain imaging data to predict non-brain-imaging phenotypes in individual participants. However, most prediction studies are underpowered, relying on less than a few hundred participants, leading to low reliability and inflated prediction performance. Yet, small sample sizes are unavoidable when studying clinical populations or addressing focused neuroscience questions. Here, we propose a simple framework – “meta-matching” – to translate predictive models from large-scale datasets to *new unseen* non-brain-imaging phenotypes in boutique studies. The key observation is that many large-scale datasets collect a wide range inter-correlated phenotypic measures. Therefore, a unique phenotype from a boutique study likely correlates with (but is not the same as) some phenotypes in some large-scale datasets. Meta-matching exploits these correlations to boost prediction in the boutique study. We applied meta-matching to the problem of predicting non-brain-imaging phenotypes using resting-state functional connectivity (RSFC). Using the UK Biobank (N = 36,848), we demonstrated that meta-matching can boost the prediction of new phenotypes in small independent datasets by 100% to 400% in many scenarios. When considering relative prediction performance, meta-matching significantly improved phenotypic prediction even in samples with 10 participants. When considering absolute prediction performance, meta-matching significantly improved phenotypic prediction when there were least 50 participants. With a growing number of large-scale population-level datasets collecting an increasing number of phenotypic measures, our results represent a lower bound on the potential of meta-matching to elevate small-scale boutique studies.

## 1. Introduction

Individual-level prediction is a fundamental goal in systems neuroscience and is important for precision medicine (Gabrieli et al., 2015; Woo et al., 2017; Eickhoff and Langner, 2019; Varoquaux and Poldrack, 2019). Therefore, there is growing interest in leveraging brain imaging data to predict non-brain-imaging phenotypes (e.g., fluid intelligence or clinical outcomes) in individual participants. To date however, most prediction studies are underpowered, including less than a few hundred participants. This has led to systemic issues related to low reproducibility and inflated prediction performance (Arbabshirani et al., 2017; Bzdok and Meyer-Lindenberg, 2018; Masouleh et al., 2019; Poldrack et al., 2020). Prediction performance can dramatically improve when training models with well-powered samples (Chu et al., 2012; Cui and Gong, 2018; He et al., 2020; Schulz et al., 2020). The advent of large-scale population-level human neuroscience datasets (e.g., UK Biobank, ABCD) is therefore critical to improving the performance and reproducibility of individual-level prediction. However, when studying clinical populations or addressing focused neuroscience topics, small-scale datasets are unavoidable. Here, we proposed a simple framework to effectively translate predictive models from large-scale datasets to *new* non-brain-imaging phenotypes (shortened as phenotypes) in small data.

More specifically, given a large-scale brain imaging dataset (N > 10,000) with multiple phenotypes, we seek to translate models trained from the large dataset to new unseen phenotypes in a small independent dataset (N ≤ 200). We emphasize that the large and small datasets are independent. Furthermore, phenotypes in the small independent dataset do not have to overlap with those in the large dataset. In machine learning, this problem is known as meta-learning or learning-to-learn (Andrychowicz et al., 2016; Finn et al., 2017; Ravi and Larochelle, 2017; Vanschoren, 2019). For example, meta-learning can be applied to a large dataset (e.g., one million natural images) to train a deep neural network (DNN) to recognize multiple object categories (e.g., furniture and humans). The DNN can then be adapted to recognize a *new* unseen object category (e.g., birds) with few samples (Nichol et al., 2018; X. Li et al., 2019; Mahajan et al., 2020). By learning a common representation across many object categories, meta-learning is able to adapt the DNN to a new object category with relatively few examples (Rusu et al., 2019; X. Li et al., 2019; Mahajan et al., 2020).

The key observation underpinning our meta-learning approach is that the vast majority of phenotypes are not independent, but are inter-correlated (Figure 1). Indeed, previous studies have discovered a relatively small number of components linking brain imaging data and an entire host of non-brain-imaging phenotypes, such as, cognition, mental health, demographics and other health attributes (Smith et al., 2015; Miller et al., 2016; Alnæs et al., 2020). Therefore, a unique phenotype X examined by a small-scale boutique study is probably correlated with (but not the same as) a particular phenotype Y in some preexisting large-scale population dataset. Consequently, a machine learning model that has been trained on phenotype Y in the large-scale dataset might be readily translated to phenotype X in the boutique study. In other words, meta-learning can be instantiated in human neuroscience by exploiting this existing correlation structure, a process we refer to as “meta-matching”.

**Figure 1.**
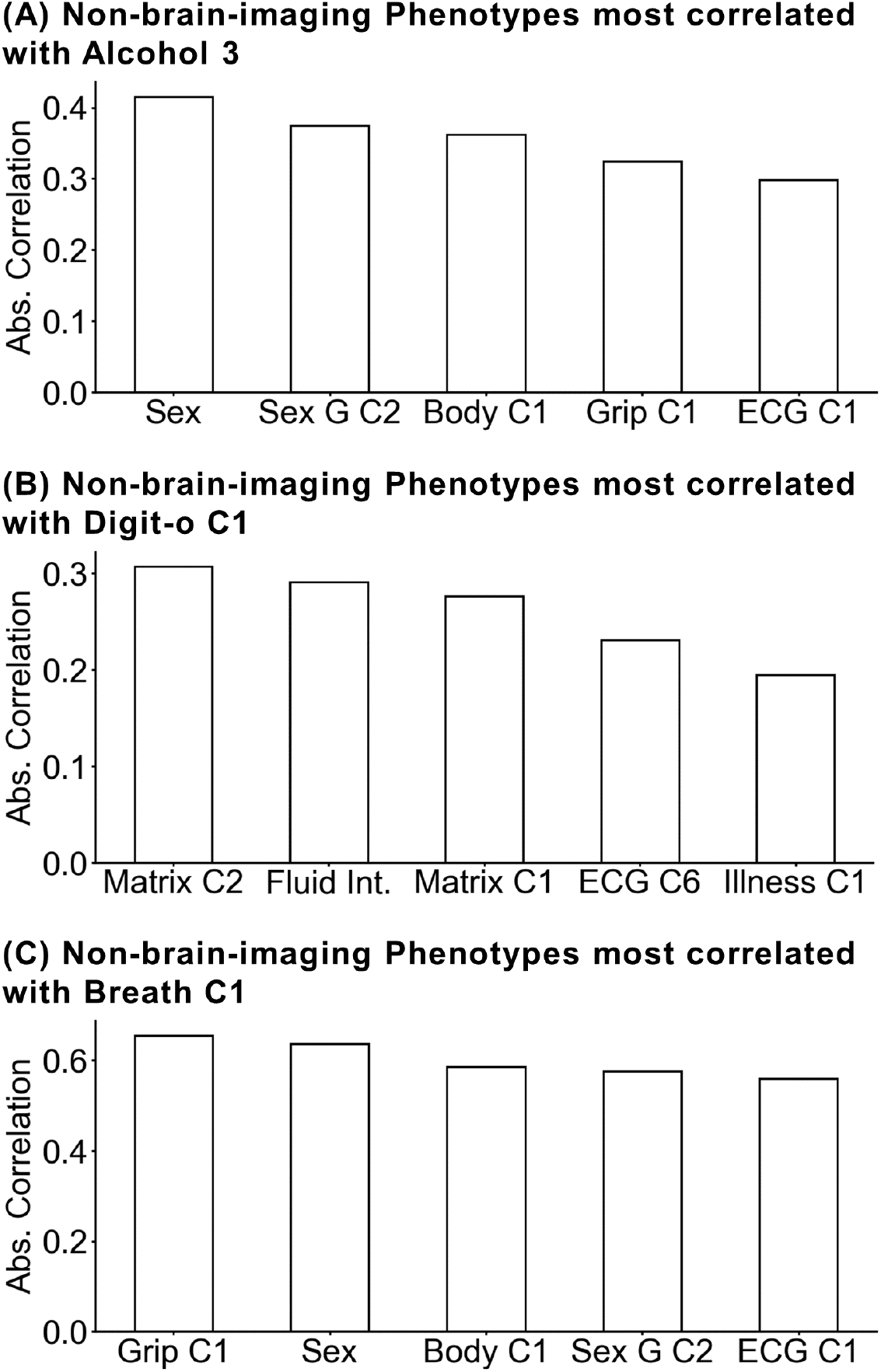
Non-brain-imaging phenotypes are inter-correlated. Here, we show the top top five absolute Pearson’s correlations of three phenotypes in the UK Biobank. (A) Top five phenotypes correlated with “Alcohol 3” (average weekly beer plus cider intake) (B) Top five phenotypes correlated with “Digit-o C1” (symbol digit substitution online principal component 1) (C) Top five phenotypes correlated with “Breath C1” (spirometry principal component 1). In selecting these examples, “Alcohol 3”, “Digit-o C1” and “Breath C1” were selected from the test meta-set, while the top five correlated phenotypes were constrained to be from the training meta-set (Figure 2). “Sex G C2” denotes genotype sex inference principal component 2. “Body C1” denotes anthropometry principal component 1. “Grip C1” denotes hand grip strength principal component 1. “ECG C1” denotes ECG measures principal component 1. “Matrix C2” denotes matrix pattern completion principal component 2. “Fluid Int.” denotes fluid intelligence. “Matrix C1” denotes matrix pattern completion principal component 1. “ECG C6” denotes ECG measures principal component 6. “Illness C1” denotes non-cancer illness principal component 1.

Meta-matching can be broadly applied to different types of magnetic resonance imaging (MRI) data. Here, we focused on the use of resting-state functional connectivity (RSFC) to predict non-brain-imaging phenotypes. RSFC measures the synchrony of resting-state functional MRI signals between brain regions (Biswal et al., 1995; Fox and Raichle, 2007; Buckner et al., 2013), while participants lie at rest without any “extrinsic” task. RSFC has provided important insights into human brain organization across health and disease (Smith et al., 2009; Yeo et al., 2011; Fornito et al., 2015; Xia et al., 2018; Kebets et al., 2019). Given any brain parcellation atlas (e.g., Shen et al., 2013; Glasser et al., 2016; Gordon et al., 2016; Eickhoff et al., 2018), a whole brain RSFC matrix can be computed for each participant. Each entry in the RSFC matrix reflects the functional coupling strength between two brain parcels. In recent years, there is increasing interest in the use of RSFC for predicting non-brain-imaging phenotypes (e.g., age or cognition) of individual participants, i.e., functional connectivity fingerprint (Dosenbach et al., 2010; Finn et al., 2015; Rosenberg et al., 2016; Reinen et al., 2018; J. Li et al., 2019; Weis et al., 2020). Thus, our study will utilize RSFC-based phenotypic prediction to illustrate the power and field-wide utility of meta-matching.

To summarize, we proposed meta-matching, a simple framework to exploit large-scale brain imaging datasets for boosting RSFC-based prediction of new, unseen phenotypes in small datasets. The meta-matching framework is highly flexible and can be coupled with any machine learning algorithm. Here, we considered kernel ridge regression (KRR) and fully connected deep neural network (DNN), which we have previously demonstrated to work well for RSFC-based behavioral and demographics prediction (He et al., 2020). We developed two classes of meta-matching algorithms: basic and advanced. Evaluation of our meta-matching approach using almost 40,000 participants from the UK Biobank (Sudlow et al., 2015; Miller et al., 2016) suggests that meta-matching can outperform classical approaches with 100% to 400% improvement in most scenarios.

## 2. Results

### 2.1 Experimental setup and classical machine learning baseline

Our study utilized 55 x 55 resting-state functional connectivity (RSFC) matrices from 36,848 participants and 67 non-brain-imaging phenotypes from the UK Biobank (Sudlow et al., 2015). The 67 phenotypes were winnowed down from an initial list of 3,937 phenotypes by a systematic procedure that excluded brain variables, binary variables (except sex), repeated measures and measures missing from too many participants. Phenotypes that were not predictable even with 1000 participants were also excluded; note that these 1000 participants were excluded from the 36,848 participants. See Section 4.4 for more details.

The data was randomly divided into training (N = 26,848; 33 phenotypes) and test (N = 10,000; 34 phenotypes) meta-sets (Figure 2A). No participant or phenotype overlapped across the training and test meta-sets. Figure 2B shows the absolute Pearson’s correlations between the training and test phenotypes. The test meta-set was further split into K participants (K-shot; K = 10, 20, 50, 100, 200) and remaining 10,000-K participants. The group of K participants served to mimic traditional small-N studies.

**Figure 2.**
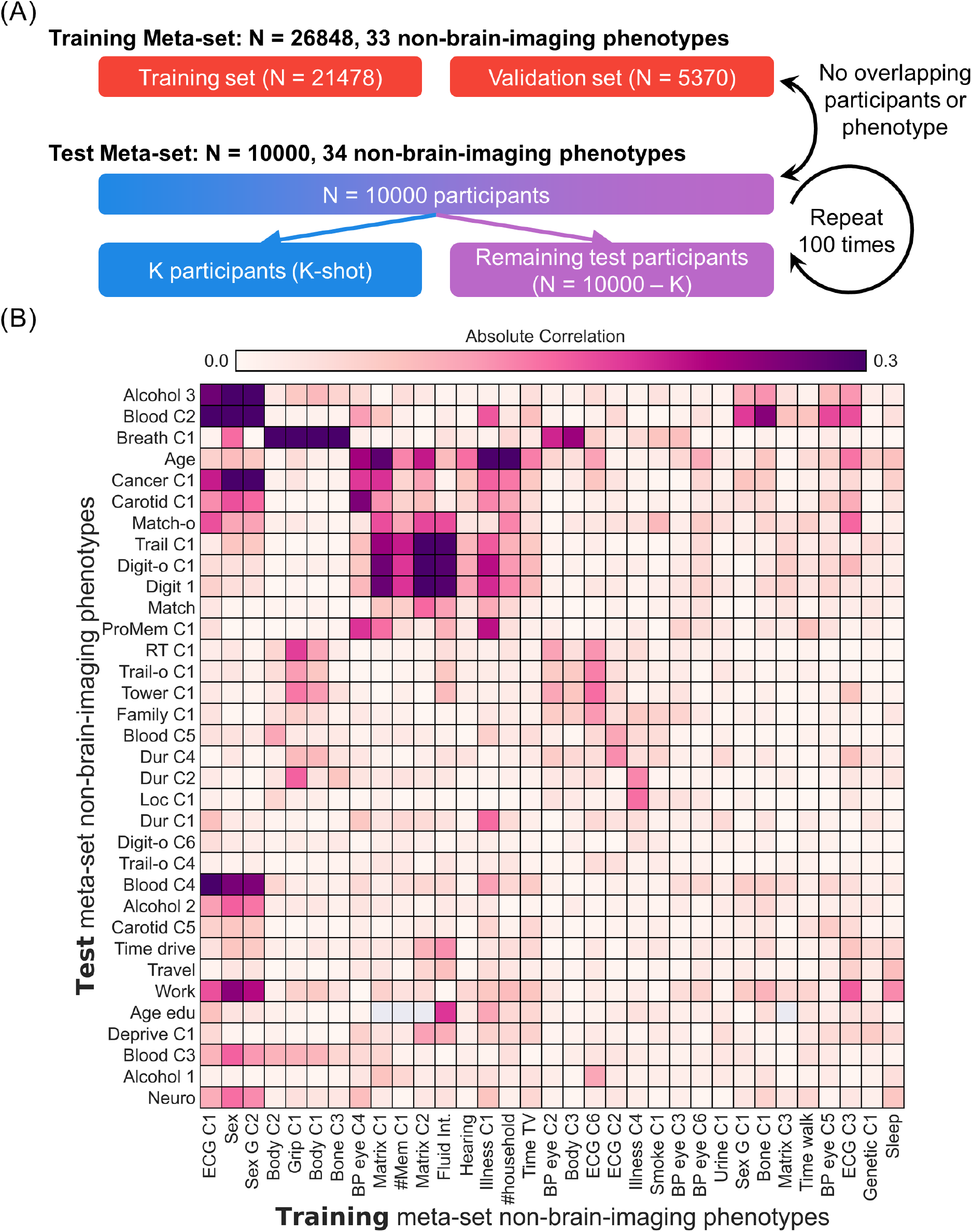
Experimental setup for meta-matching. The goal of meta-matching is to translate predictive models from big datasets to new unseen non-brain-imaging phenotypes in independent small datasets. (A) UK Biobank dataset (Jan 2020 release) was divided into a training meta-set comprising 26,848 participants and 33 non-brain-imaging phenotypes, and a test meta-set comprising independent 10,000 participants and 34 other phenotypes. It is important to emphasize that no participant and phenotype overlapped between training and test meta-sets. The test meta-set was in turn split into K participants (K = 10, 20, 50, 100, 200) and remaining 10,000-K participants. The group of K participants mimicked studies with traditionally common sample sizes. This split was repeated 100 times for robustness. (B) Absolute Pearson’s correlations between phenotypes in training and test metasets. Each row represents one test meta-set phenotype. Each column represents one training meta-set phenotype. Figures S1 and S2 show correlation plots for phenotypes within training and test meta-sets. Dictionary of phenotypes is found in Tables S1 and S2.

For each phenotype in the test meta-set, a classical machine learning baseline (kernel ridge regression; KRR) was trained on the RSFC matrices of the K participants and applied to the remaining 10,000 – K participants. Hyperparameters were tuned on the K participants. We note that small-N studies obviously do not have access to the remaining 10,000 – K participants. However, in our experiments, we utilized a large sample of participants (10,000 – K), so as to accurately establish the performance of the classical machine learning baseline. We repeated this procedure 100 times (each with different sample of K participants) to ensure robustness of the results (Varoquaux et al., 2017).

KRR was chosen as a baseline because of the small number of hyperparameters, which made it suitable for small-N studies. We have also previously demonstrated that KRR and deep neural networks can achieve comparable prediction performance in functional connectivity prediction of behavior and demographics in both small-scale and large-scale datasets (He et al., 2020).

### 2.2 Basic meta-matching dramatically outperforms classical machine learning baseline

The meta-matching framework is highly flexible and can be instantiated with different machine learning algorithms. Here, we considered kernel ridge regression (KRR) and fully connected deep neural network (DNN), which we have previously demonstrated to work well for RSFC-based behavioral and demographics prediction (He et al., 2020). We considered two classes of meta-matching algorithms: basic and advanced (Figure 3).

**Figure 3.**
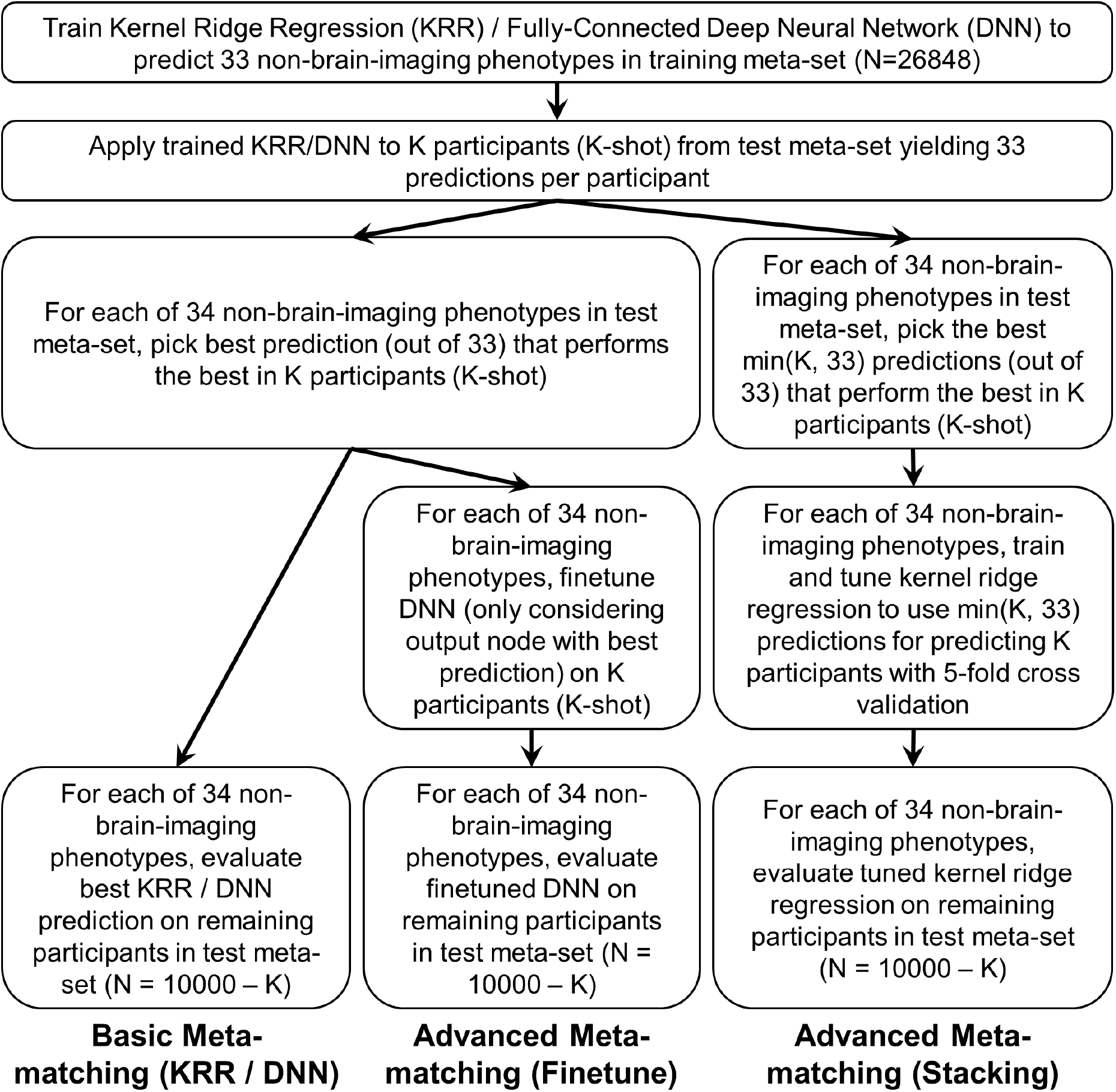
Application of basic and advanced meta-matching to the UK Biobank. The meta-matching framework can be instantiated using different machine learning algorithms. Here, we incorporated kernel ridge regression (KRR) and fully-connected feedforward deep neural network (DNN) within the meta-matching framework. We proposed two classes of meta-matching algorithms: basic and advanced. In the case of basic meta-matching, we considered two variants: basic meta-matching (KRR) and basic meta-matching (DNN). In the case of advanced meta-matching, we considered two variants: advanced meta-matching (finetune) and advanced meta-matching (stacking). Both advanced meta-matching variants utilized the DNN. See text for more details.

In “basic meta-matching (KRR)”, for each non-brain-imaging phenotype in the training meta-set, we trained a kernel ridge regression (KRR) model to predict the phenotype from the RSFC matrices. We then applied the 33 trained KRR models to the RSFC of the K participants (from the test meta-set), yielding 33 predictions per participant. For each test meta-set phenotype, we picked the prediction (out of 33 predictions) that predicted the test meta-set phenotype the best in the K participants. The corresponding KRR model (yielding this best prediction) was used to predict the test phenotype in the remaining 10,000 – K participants. We also repeated the above procedure using a generic fully connected feedforward deep neural network (DNN) instead of KRR, yielding the “basic meta-matching (DNN)” algorithm. The only difference is that instead of training 33 DNNs (which would require too much computational time), a single 33-output DNN was utilized. See Section 4.7 for details.

Figure 4A shows the prediction accuracies (Pearson’s correlation coefficient) averaged across 34 phenotypes and 10,000 – K participants in the test meta-set. The boxplots represent 100 random repeats of K participants (K-shot). Bootstrapping was utilized to derive p values (Figure 4B; Figure S3; see Methods). Multiple comparisons were corrected using false discovery rate (FDR, q < 0.05). Both basic meta-matching algorithms were significantly better than the classical (KRR) approach across all sample sizes (Figure 4B). The improvements were large. For example, in the case of 20-shot (a typical sample size for many fMRI studies), basic meta-matching (DNN) was more than 100% better than classical (KRR): 0.124 ± 0.016 (mean ± std) versus 0.052 ± 0.007. Indeed, classical (KRR) required 200 participants before achieving an accuracy (0.120 ± 0.005) comparable to basic meta-matching (DNN) with 20 participants.

**Figure 4.**
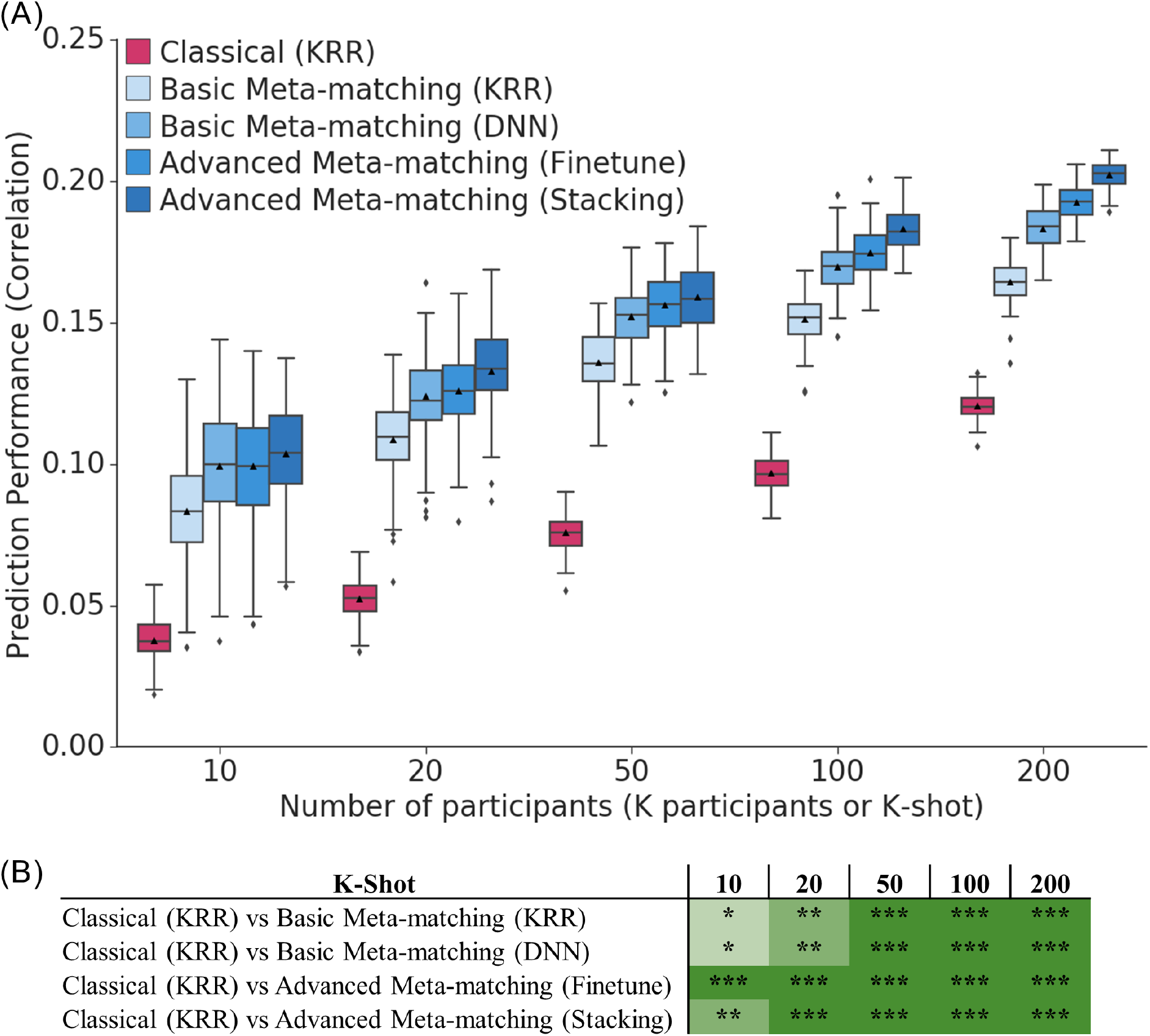
Meta-matching reliably outperforms predictions from classical kernel ridge regression (KRR). (A) Prediction performance (Pearson’s correlation) averaged across 34 non-brain-imaging phenotypes in the test meta-set (N = 10,000 – K). The K participants were used to train and tune the models (Figure 3). Boxplots represent variability across 100 random repeats of K participants (Figure 2A). Whiskers represent 1.5 inter-quartile range. (B) Statistical difference between the prediction performance (Pearson’s correlation) of classical (KRR) baseline and meta-matching algorithms. “*” indicates p < 0.05 and statistical significance after multiple comparisons correction (FDR q < 0.05). “**” indicates p < 0.001 and statistical significance after multiple comparisons correction (FDR q < 0.05). “***” indicates p < 0.00001 and statistical significance after multiple comparisons correction (FDR q < 0.05). The actual p values and statistical comparisons among all algorithms are found in Figure S3. Prediction performance measured using coefficient of determination (COD) is found in Figure S4.

When utilizing coefficient of determinant (COD) as a metric of prediction performance (Figures S4 and S5), all algorithms performed poorly (COD ≤ 0) when there were less than 50 participants (K < 50), suggesting worse than chance prediction. When there were at least 50 participants (K ≥ 50), basic meta-matching algorithms became substantially better than classical (KRR) approach. However, the improvement was only statistically significant starting from around 100-200 participants.

To summarize, basic meta-matching performed well even with 10 participants if the goal was “relative” prediction (i.e., Pearson’s correlation; Scheinost et al., 2019). However, if the goal was “absolute” prediction (i.e., COD; Poldrack et al., 2020), then basic meta-matching required at least 100 participants to work well.

### 2.3 Advanced meta-matching provides further improvement

We have demonstrated that basic meta-matching led to significant improvement over the classical (KRR) baseline. However, in practice, there might be significant differences between the training and test meta-sets, so simply picking the best non-brain-imaging phenotypic prediction model from the training meta-set might not generalize well to the test meta-set. Thus, we proposed two additional meta-matching approaches: “advanced meta-matching (finetune)” and “advanced meta-matching (stacking)”.

As illustrated in Figure 3, the procedure for advanced meta-matching (finetune) is similar to basic meta-matching (DNN). Briefly, we trained a single DNN (with 33 outputs) on the training meta-set. We then applied the 33-output DNN to the K participants and picked the best DNN model for each test phenotype (out of 34 phenotypes). We then finetuned the top two layers of the DNN using the K participants before applying the finetuned model to the remaining 10,000 – K participants. See Section 4.8 for details.

In the case of advanced meta-matching (stacking), we trained a single DNN (with 33 outputs) on the training meta-set. We then applied the 33-output DNN to the K participants, yielding 33 predictions per participant. The top M predictions are then used as features for predicting the phenotype of interest in the K participants using KRR. To reduce overfitting, M is set to be the minimum of 33 and K. For example, for the 10-shot scenario, M is set to be 10. For the 50-shot scenario, M is set to be 33. The DNN (which was trained on the training meta-set) and KRR models (which were trained on the K participants) were then applied to the remaining 10,000 – K participants. See Section 4.8 for details.

Figure 4A shows the prediction accuracies (Pearson’s correlation coefficient) averaged across 34 phenotypes and 10,000 – K participants in the test meta-set. Both advanced meta-matching algorithms exhibited large and statistically significant improvements over classical (KRR) approach across all sample sizes (Figure 4B). For example, in the case of 20-shot, advanced meta-matching (stacking) was more than 100% better than classical (KRR): 0.133 ± 0.014 (mean ± std) versus 0.053 ± 0.007. Among the meta-matching algorithms, the advanced meta-matching algorithms were numerically better than the basic meta-matching algorithms from 20-shot onwards, but statistical significance was not achieved until around 100-shot onwards (Figure S3B).

When utilizing coefficient of determinant (COD) as a metric of prediction performance (Figures S4 and S5), all algorithms performed poorly (COD ≤ 0) when there were less than 50 participants (K < 50), suggesting chance or worse than chance prediction. From 50-shot onwards, advanced meta-matching algorithms became statistically better than classical (KRR) approach (Figures S4B and S5B). The improvements were substantial. For example, in the case of 100-shot, advanced meta-matching (stacking) was 400% better than classical (KRR): 0.053 ± 0.005 (mean ± std) versus 0.010 ± 0.004. Among the meta-matching algorithms, the advanced meta-matching algorithms were numerically better than the basic meta-matching algorithms from 100-shot onwards, but statistical significance was not achieved until 200-shot (Figure S5B).

To summarize, advanced meta-matching performed well even with 10 participants if the goal was “relative” prediction (i.e., Pearson’s correlation; Scheinost et al., 2019). However, if the goal was “absolute” prediction (i.e., COD; Poldrack et al., 2020), then advanced meta-matching required at least 50 participants to work well.

### 2.4 Prediction improvements were driven by correlations between training and test meta-set non-brain-imaging phenotypes

Figures 5 illustrates the 100-shot prediction performance (Pearson’s correlation coefficient) of four test meta-set non-brain-imaging phenotypes across all approaches. Figure S6 shows the same plot for COD. For three of the phenotypes (average weekly beer plus cider intake, symbol digit substitution and matrix pattern completion), meta-matching demonstrated substantial improvements over classical (KRR). In the case of the last phenotype (time spent driving per day), meta-matching did not yield any statistically significant improvement.

**Figure 5.**
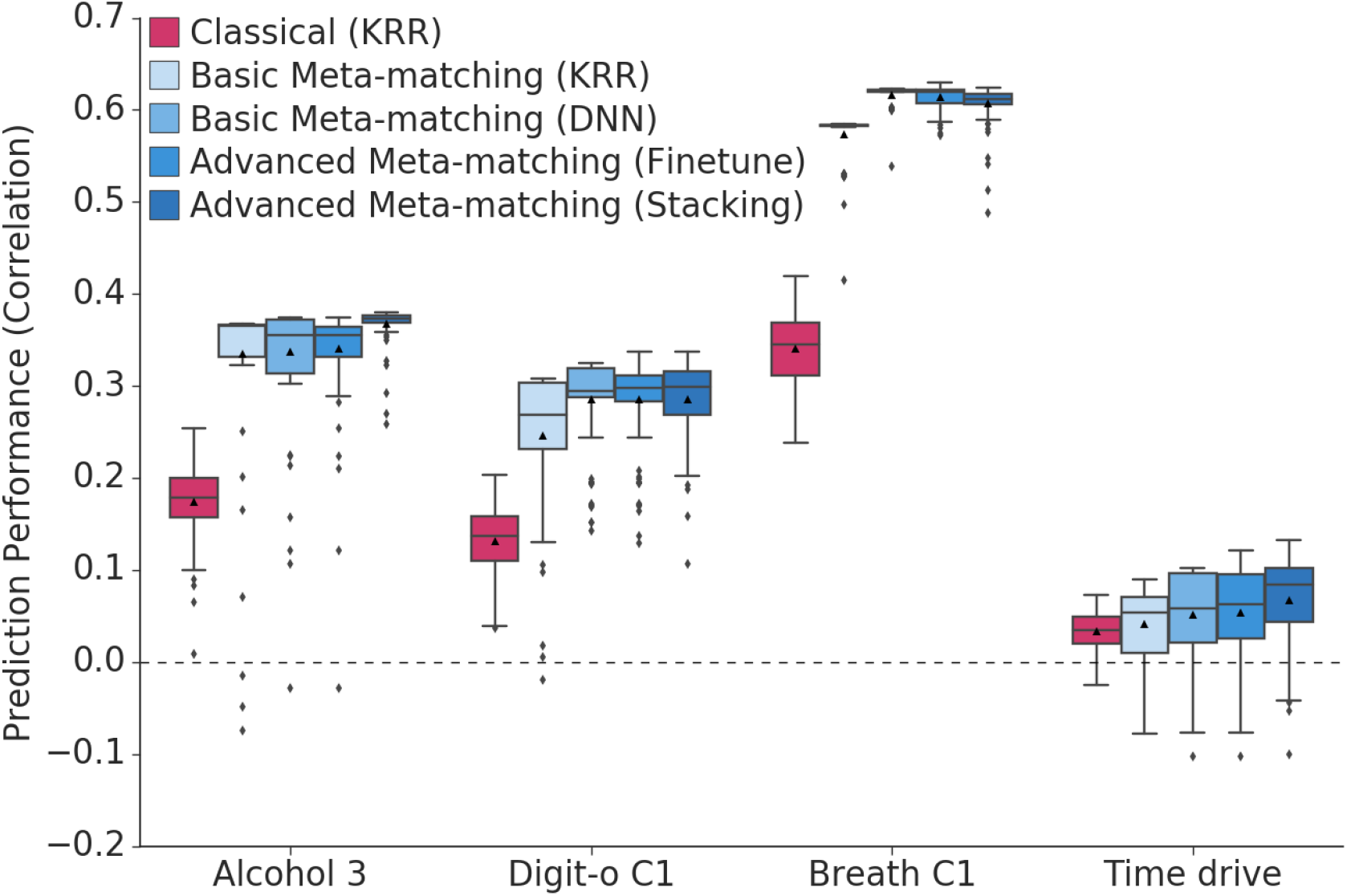
Examples of non-brain-imaging phenotypic prediction performance in the test meta-set in the case of 100-shot learning. Here, prediction performance was measured using Pearson’s correlation. “Alcohol 3” (average weekly beer plus cider intake) was most frequently matched to “Bone C3” (bone-densitometry of heel principal component 3). “Digit-o C1” (symbol digit substitution online principal component 1) was most frequently matched to “Matrix C1” (matrix pattern completion principal component 1). “Breath C1” (spirometry principal component 1) was most frequently matched to “Grip C1” (hand grip strength principal component 1). “Time drive” (Time spent driving per day) was most frequently matched to “BP eye C3” (blood pressure & eye measures principal component 3). Figure S6 shows an equivalent figure using coefficient of determination (COD) as the prediction performance measure.

Given that meta-matching exploits correlations among phenotypes, we hypothesized that variability in prediction improvements were driven by inter-phenotype correlations between the training and test meta-sets. Figure 6 shows the performance improvement (Pearson’s correlation) as a function of the maximum correlation between each test phenotype and training phenotypes. Figure S7 shows the same plot for COD. As expected, test phenotypes with stronger correlations with at least one training phenotype led to greater prediction improvement with meta-matching. Interestingly, phenotypes that were better predicted by classical (KRR) also benefited more from meta-matching (Figure S8).

**Figure 6.**
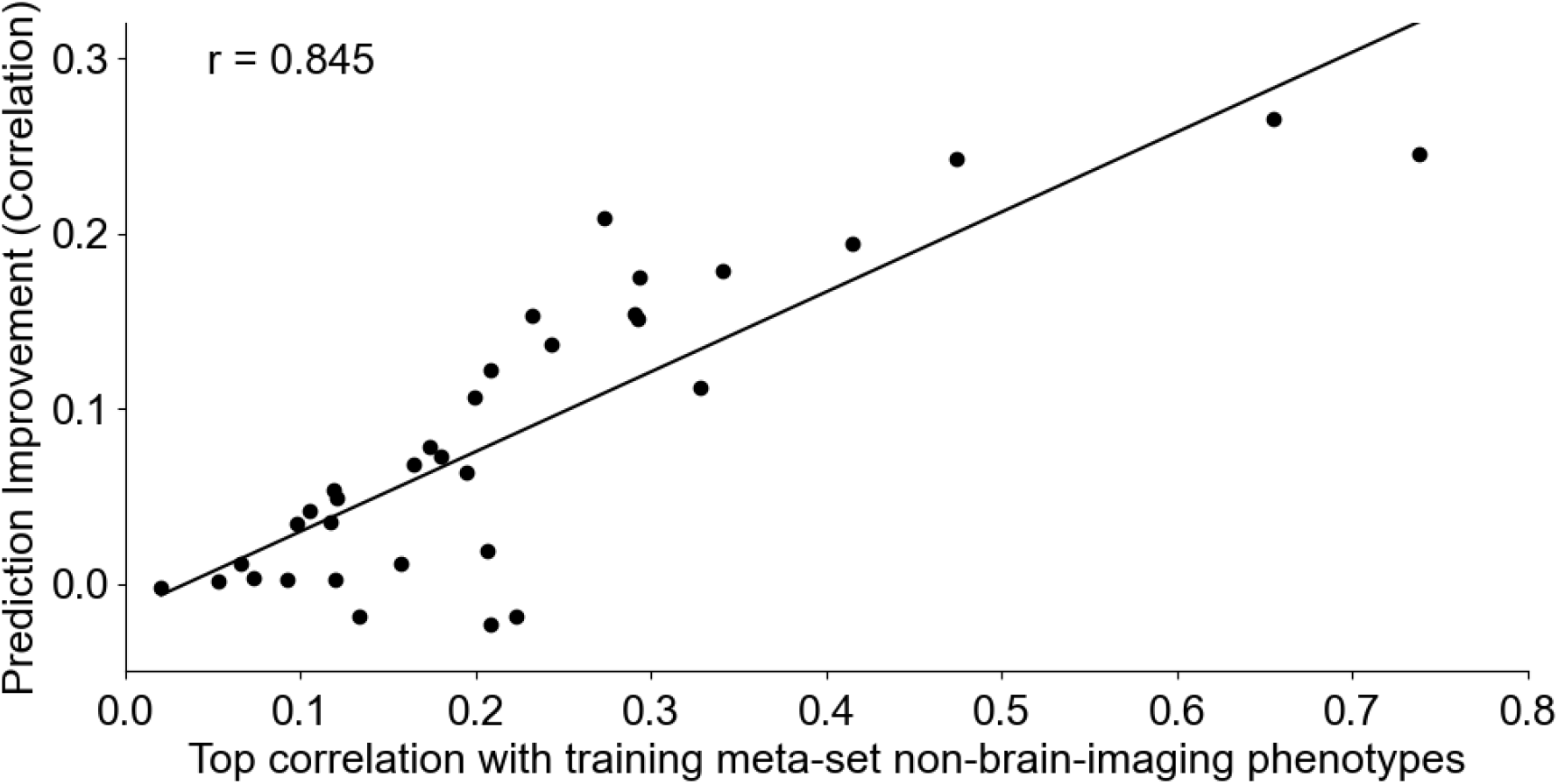
Prediction improvements were driven by correlations between training and test meta-set phenotypes. Vertical axis shows the prediction improvement of advanced meta-matching (stacking) with respect to classical (KRR) baseline under the 100-shot scenario. Prediction performance was measured using Pearson’s correlation. Each dot represents a test meta-set phenotype. Horizontal axis shows each test phenotype’s top absolute Pearson’s correlation with phenotypes in the training meta-set. Test phenotypes with stronger correlations with at least one training phenotype led to greater prediction improvement with meta-matching. Similar conclusions were obtained with coefficient of determination (Figure S7).

### 2.5 Computational costs of meta-matching

Meta-matching comprises two stages: training on the training meta-set and meta-matching on new non-brain-imaging phenotypes in the K participants (K-shot). Training and hyperparameter tuning on the training meta-set is slow, but only has to be performed once. For example, in our study, training the DNN with automatic hyperparameter tuning using the HORD algorithm (Regis and Shoemaker, 2013; Ilievski et al., 2017; Eriksson et al., 2019) on a single graphics processing unit (GPU) took about 2 days. In the case of both basic meta-matching algorithms, meta-matching on new non-brain-imaging phenotypes is extremely fast because it only requires forward passes through a neural network (in the case of DNN) or matrix multiplications (in the case of KRR). More specifically, the second stage for basic meta-matching algorithms took less than 0.1 second for a single test meta-set phenotype and one K-shot. In the case of advanced meta-matching (stacking), there is an additional step of training a KRR model on the K participants. Nevertheless, the second stage for advanced meta-matching (stacking) only took 0.5 second for a single meta-set phenotype and one K-shot. On the other hand, the computational cost for finetuning the DNN for advanced meta-matching (finetune) is a lot more substantial, requiring about ~30 seconds for a single test meta-set phenotype and one K-shot. Although 30 seconds might seem quite fast, repeating the K-shot 100 times for all values of K and 34 meta-set phenotypes required 6 full days of computational time.

## 3. Discussion

In this study, we proposed “meta-matching”, a simple framework to effectively translate predictive models from large-scale datasets to *new* non-brain-imaging phenotypes in small data. Using a large sample of almost 40,000 participants from the UK Biobank, we demonstrated that meta-matching can dramatically boost prediction performance in the small-sample scenario. Overall, our results suggest that meta-matching will be extremely helpful for boosting the predictive power in small-scale boutique studies focusing on specific neuroscience questions or clinical populations.

We note that a variety of prediction performance measures have been utilized in the literature. For studies interested in relative ranking (Finn et al., 2015; Scheinost et al., 2019), Pearson’s correlation is a common performance metric. We showed that if Pearson’s correlation was used as a performance metric, meta-matching performed very well even with as few as 10 participants (Figure 4). Thus, if the experimenter’s goal is relative ranking, then our experiments suggest that meta-matching is superior regardless of sample sizes.

However, others have strongly argued in favor of absolute prediction performance (Poldrack et al., 2020). In this scenario, coefficient of determination (COD) is a common performance metric. We showed that if COD was used as a performance metric, meta-matching dramatically outperformed classical (KRR) where there were more than 50 participants (Figure S4). However, when there were only 10-20 participants, meta-matching was statistically worse than classical (KRR). However, we note that in this small sample scenario, COD was less than or equal to zero even with classical (KRR). Thus, our experiments suggest that absolute prediction is unlikely to be successful in the case of 10-20 participants and should not be considered a realistic goal.

A major limitation of meta-matching is that it exploits correlations between training and test meta-sets, so the amount of prediction improvement strongly relied on the strongest correlations between the test phenotype and training phenotypes (Figure 6). However, we note that this limitation will ameliorate with increasing number of large-scale population-level datasets collecting an increasing number of phenotypes, which would increase the probability of some phenotypes in some large-scale datasets being correlated with a new phenotype of interest in a smaller dataset. Both advanced meta-matching (finetune) and advanced meta-matching (stacking) seek to address this limitation by further adaptation to the K participants in the test meta-set. However, when considering absolute prediction performance, the more advanced adaptation procedures required sufficient number of participants to work well. For example, advanced meta-matching (stacking) was statistically better than simple meta-matching (KRR) only when there were at least 50-100 participants when considering COD as a performance metric.

Finally, it might be worthwhile to differentiate our approach from transfer learning, which is a lot more common in brain imaging studies (Koppe et al., 2020). In transfer learning applications, DNNs are typically pre-trained using a large-scale dataset (source domain), which might or might not comprise brain imaging data. The pretrained DNN is then finetuned for a prediction problem in another dataset (target domain; Cheng et al., 2015; Kim et al., 2016; Heinsfeld et al., 2018; Chen et al., 2020). In transfer learning, the pretraining in the source domain typically involves no prediction problem or a problem that is very similar to the target domain. By contrast, meta-learning involves training a machine learning model from a wide range of prediction problems in the source domain, which can then be adapted to a *new* prediction problem in the target domain.

## Conclusion

In this study, we proposed a simple framework, meta-matching, that allowed the translation of phenotypic predictive models from large-scale datasets to *new* phenotypes in small-scale datasets. Using a large sample of almost 40,000 participants, we demonstrated that meta-matching could dramatically boost phenotypic prediction performance in as few as 10 participants. We believe that meta-matching will be extremely useful for small-scale boutique studies focusing on clinical populations or specific neuroscience questions.

## 4. Methods

### 4.1 Datasets

This study utilized data from the UK Biobank (Sudlow et al., 2015; Miller et al., 2016) under UK Biobank resource application 25163. The UK Biobank is a population epidemiology study with 500,000 adults (age 40-69) recruited between 2006 and 2010 (Sudlow et al., 2015). A subset of 100,000 participants is being recruited for multimodal imaging, including brain MRI, e.g., structural MRI and resting-state fMRI (rs-fMRI) from 2016 to 2022 (Miller et al., 2016). A wide range of non-brain-imaging phenotypes was acquired for each participant. Here, we considered the January 2020 release of 37,848 participants with structural MRI and rs-fMRI. Structural MRI (1.0mm isotropic) and rs-fMRI (2.4mm isotropic) were acquired at four imaging centers (Bristol, Cheadle Manchester, Newcastle, and Reading) with harmonized Siemens 3T Skyra MRI scanners. Each subject has one rs-fMRI run with 490 frames (6 minutes) and a TR of 0.735s. More details of the UK Biobank dataset can be found elsewhere (Elliott and Peakman, 2008; Sudlow et al., 2015; Miller et al., 2016; Alfaro-Almagro et al., 2018).

### 4.2 Brain imaging data

Our study utilized the 55 x 55 RSFC (partial correlation, Smith et al., 2011) matrices from data-field 25753 of the UK Biobank (Miller et al., 2016; Alfaro-Almagro et al., 2018). Details about rs-fMRI preprocessing can be found elsewhere (Alfaro-Almagro et al., 2018). Data-field 25753 RSFC had 100 whole-brain spatial independent component analysis (ICA) derived components (Beckmann and Smith, 2004). After the removal of 45 artifactual components, as indicated by the UKB team, 55 components were left (Miller et al., 2016). Data-field 25753 contains two instances, first imaging visit (instance 2) and first repeat imaging visit (instance 3). The first imaging visit (instance 2) had RSFC data for 37,848 participants, while the first repeat imaging visit (instance 3) only had RSFC data for 1,493 participants. Here, we only considered RSFC from the first imaging visit (instance 2).

### 4.3 RSFC-based prediction setup

Our meta-matching framework is highly flexible and can be instantiated with different machine learning algorithms. Here, we considered kernel ridge regression (KRR) and fully-connected feedforward deep neural network (DNN), which we have previously demonstrated to work well for RSFC-based behavioral and demographics prediction (He et al., 2020). As discussed in the previous section, each RSFC matrix was a symmetric 55 x 55 matrix, where 55 is the number of independent components. Each element represented the degree of statistical dependencies between two brain components. The lower triangular elements of the RSFC matrix of each participant were then vectorized and used as input features for KRR and DNN to predict individuals’ phenotypes.

Kernel ridge regression (Murphy, 2012) is a non-parametric machine learning algorithm. This method is a natural choice as we previously demonstrated that KRR achieved similar prediction performance as several deep neural networks (DNNs) for the goal of RSFC-based behavioral and demographics prediction (He et al., 2020). Roughly speaking, KRR predicts the phenotype (e.g., fluid intelligence) of a test participant by the weighted average of all training participants’ phenotypes (e.g., fluid intelligence). The weights in the weighted average is determined by the similarity (i.e., kernel) between the test participant and training participants. In this study, similarity between two participants was defined as the Pearson’s correlation between the vectorized lower triangular elements of their RSFC matrices. KRR also contains an *l*_2_ regularization term as part of the loss function to reduce overfitting. The hyperparameter λ is used to control the strength of the *l*_2_ regularization. More details can be found elsewhere (Murphy, 2012; He et al., 2020).

A fully-connected feedforward deep neural network (DNN) is one of the most classical DNNs (Lecun et al., 2015). We previously demonstrated that the feedforward DNN and KRR could achieve similar performance for RSFC-based behavioral and demographics prediction (He et al., 2020). In this study, the DNN was trained based on the vectorized lower triangular elements of the RSFC matrix as input features and output the prediction of one or more non-brain-imaging phenotypes. The DNN consists of several fully connected layers. Each node (except input layer nodes) is connected to all nodes in the previous layer. The values at each node is the weighted sum of node values from the previous layer. The outputs of the hidden layer nodes go through a nonlinear activation function, Rectified Linear Units (ReLU; *f*(*x*) = *max*(0, *x*)). The output layer is linear. More details about hyperparameter tuning (e.g., number of layers and number of nodes per layer) are found in Supplemental Methods S1.

### 4.4 Non-brain-imaging phenotype selection and processing

To obtain the final set of 67 non-brain-imaging phenotypes, we began by extracting all 3,937 unique phenotypes available to us under UK Biobank resource application 25163. We then performed three stages of selection and processing:

1. In the first stage, we

- Removed non-continuous and non-integer data fields (date and time converted to float), except for sex.
- Removed Brain MRI phenotypes (category ID 100).
- Removed first repeat imaging visit (instance 3).
- Removed first two instances (instance 0 and 1) if first imaging visit (instance 2) exists and first imaging visit (instance 2) participants were more than double of participants from instance 0 or 1.
- Removed first instance (instance 0) if only the first two instances (instance 0 and 1) exist, and instance 1 participants were more than double of participants from instance 0.
- Removed phenotypes for which less than 2000 participants had RSFC data.
- Removed behaviors with the same value for more than 80% of participants. After the first stage of filtering, we were left with 701 phenotypes.
2. We should not expect every phenotype to be predictable by RSFC. Therefore, in the second stage, our goal was to remove phenotypes that could not be well predicted even with large number of participants. More specifically,

- We randomly selected 1000 participants from 37,848 participants. These 1000 participants were completely excluded from the main experiments (Figure 2A).
- Using these 1000 participants, Kernel ridge regression (KRR) was utilized to predict each of the 701 phenotypes using RSFC. To ensure robustness, we performed 100 random repeats of training, validation, and testing (60%, 20%, and 20%). For each repeat, KRR was trained on the training set and hyperparameters were tuned on the validation set. We then evaluated the trained KRR on the test set. phenotypes with an average test prediction performance (Pearson’s correlation) less than 0.1 were removed. At the end of this second stage, 265 phenotypes were left.
3. Many of the remaining phenotypes were highly correlated. For example, the bone density measurements of different body parts were highly correlated. Principal component analysis (PCA) was performed separately on each subgroup of highly similar phenotypes in the 1000-participant sample. Similarity was evaluated based on the UK Biobank-provided categories of item sets (i.e., items under the same category were considered highly similar). PCAs were not applied to 18 phenotypes (out of 265 phenotypes), which were not similar to other phenotypes. For the purpose of carrying out PCA, missing values were filled in with the EM algorithm (Seabold and Perktold, 2010). For each PCA, we kept enough components to explain 95% of the variance in the data or 6 components, whichever is lower. Overall, the PCA step reduced the 247 phenotypes (out of 265 phenotypes) to 93 phenotypes. We then repeated the previous step (stage 2) on these 93 phenotypes, resulting in 49 phenotypes with prediction performance (Pearson’s correlation) larger than 0.1. Adding back the 18 phenotypes that were not processed by PCA, we ended up with 67 phenotypes utilized in the main experiment (Figure 2A). This PCA procedure was also applied separately to the training and test meta-sets defined in the next section. Final list of the phenotypes is found in Tables S1 and S2.

### 4.5 Data split scheme

After the non-brain-imaging phenotypes selection process in the previous section, we were left with 36,848 participants with RSFC matrices and 67 phenotypes. As illustrated in Figure 2A, we randomly split the data into two meta-sets: training meta-set with 26,848 participants and 33 phenotypes, and test meta-set with 10,000 participants and 34 phenotype. There was no overlap between the participants and phenotypes across the two meta-set. Figure 2B shows the Pearson’s correlations between the training and test phenotypes. Figures S1 and S2 show correlation plots for phenotypes within training and test meta-sets.

For the training meta-set, we further randomly split it into a training set with 21,478 participants (80% of 26,848 participants) and validation set with 5370 participants (20% of 26,848 participants). For the test meta-set, we randomly split 10,000 participants into K participants (K-shot) and 10,000 – K participants, where K had a value of 10, 20, 50, 100, and 200. The group of K participants mimicked traditional small-N studies. Each random K-shot split was repeated 100 times to ensure stability.

Z-normalization (transforming each variable to have zero mean and unit variance) was applied to the phenotypes. In the case of the training meta-set, z-normalization was performed by using the mean and standard deviation computed from the training set within the training meta-set. In the case of the test meta-set, for each of the 100 repeats of the K-shot learning, the mean and standard deviation were computed from the K participants and subsequently applied to the full test meta-set.

### 4.6 Classical Kernel Ridge Regression (KRR) Baseline

For the classical (KRR) baseline, we performed K-shot learning for each non-brainimaging phenotype in the test meta-set, using K participants from the random split (Figure 2A). More specifically, for each phenotype, we performed 5-fold cross-validation on the K participants using different values of the hyperparameter λ (that controlled the strength of the *l*_2_ regularization). For the purpose of choosing the best hyperparameter, prediction performance was evaluated using the coefficient of determination (COD). The best hyperparameter λ was used to train the KRR model using all K participants. The trained KRR model was then applied to the remaining test participants (N = 10,000 – K). Prediction performance in the 10,000 - K participants was measured using Pearson’s correlation and COD. This procedure was repeated for each of the 100 random subsets of K participants.

### 4.7 Basic meta-matching

The meta-matching framework is highly flexible and can be instantiated with different machine learning algorithms. Here, we incorporated kernel ridge regression (KRR) and fully-connected feedforward deep neural network (DNN) within the meta-matching framework. We proposed two classes of meta-matching algorithms: basic and advanced. In the case of basic meta-matching, we considered two variants: “basic meta-matching (KRR)” and “basic meta-matching (DNN)” (Figure 3).

In the case of basic meta-matching (KRR), we first trained a KRR to predict each training non-brain-imaging phenotype from RSFC. We used the training set (N = 21478) within the training meta-set for training and validation set (N = 5370) within the training meta-set for hyperparameter tuning. The hyperparameter λ was selected via a simple grid search. There were 33 phenotypes, so we ended up with 33 trained KRR models from the training meta-set. Second, we applied the 33 trained KRR models to K participants (K-shot) from the test meta-set, yielding 33 predictions per participant. Third, for each test phenotype (out of 34 phenotypes), we picked the best KRR model (out of 33 models) that performed the best (as measured by COD) on the K participants. Finally, for each test phenotype, we applied the best KRR model to the remaining participants in the test meta-set (N = 10,000 – K). Prediction performance in the 10,000 – K participants was measured using Pearson’s correlation and COD. To ensure robustness, the K-shot procedure was repeated 100 times, each with a different set of K participants.

In the case of basic meta-matching (DNN), we first trained one single DNN to predict all 33 training phenotypes from RSFC. In other words, the DNN outputs 33 predictions simultaneously. The motivation for a single multi-output DNN is to avoid the need to train and tune 33 single-output DNNs. We used the training set (N = 21478) within the training meta-set for training and validation set (N = 5370) within the training meta-set for hyperparameter tuning. Details of the hyperparameter tuning is found in Supplementary Methods S1. Second, we applied the trained DNN to the K participants (K-shot) from the test meta-set, yielding 33 different phenotypical predictions for each given participant. Third, for each test phenotype (out of 34 phenotypes), we picked the best output DNN node (out of 33 output nodes) that generated the best prediction (as measured by COD) for the K participants. Finally, for each test phenotype, we applied the predictions from the best DNN output node on the remaining 10,000 – K participants in the test meta-set. Prediction performance in the 10,000 – K participants was measured using Pearson’s correlation and COD. To ensure robustness, the K-shot procedure was repeated 100 times, each with a different set of K participants.

### 4.8 Advanced meta-matching

There might be significant differences between the training and test meta-sets. Therefore, simply picking the best non-brain-imaging phenotypic prediction model estimated from the training meta-set might not generalize well to the test meta-set. Thus, we proposed two additional meta-matching approaches: “advanced meta-matching (finetune)” and “advanced meta-matching (stacking)” (Figure 2).

In the case of advanced meta-matching (finetune), we used the same multi-output DNN from basic meta-matching (DNN). Like before, for each test phenotype (out of 34 phenotypes), we picked the best output DNN node (out of 33 output nodes) that generated the best prediction (as measured by COD) for the K participants. We retained this best output node (while removing the remaining 32 nodes) and finetuned the DNN using the K participants (K-shot). More specifically, the K participants were randomly divided into training and validation sets using a 4:1 ratio. The training set was used to finetune the weights of the last two layers of the DNN, while the remaining weights were frozen. The validation set was used to determine the stopping criterion (in terms of the number of training epochs). The finetuned DNN was applied to the remaining 10,000 – K participants in the test meta-set. We note that the finetuning procedure was repeated separately for each of 33 test phenotypes. Prediction performance in the 10,000 – K participants was measured using Pearson’s correlation and COD. To ensure robustness, the K-shot procedure was repeated 100 times, each with a different set of K participants. More details about the finetuning procedure can be found in Supplementary Methods S2.

In the case of advanced meta-matching (stacking), we used the same multi-output DNN from basic meta-matching (DNN). The DNN was applied to the K participants (K-shot) from the test meta-set, yielding 33 predictions per participant. For each test phenotype (out of 34 phenotypes), the best M predictions (as measured by COD) were selected. To reduce overfitting, M was set to be the minimum of K and 33. Thus, if K was smaller than 33, we considered the top K outputs from the multi-output DNN. If K was larger than 33, we considered all 33 outputs of the multi-output DNN. We then trained a kernel ridge regression (KRR) model using the M DNN outputs to predict the phenotype of interest in the K participants. The hyperparameter λ was tuned using grid search and 5-fold cross-validation on the K participants. The optimal λ was then used to train a final KRR model using all K participants. Finally, the KRR model was applied to the remaining 10,000 – K participants in the test meta-set. We note that this “stacking” procedure was repeated separately for each of 33 test phenotypes. Prediction performance in the 10,000 – K participants was measured using Pearson’s correlation and COD. To ensure robustness, the K-shot procedure was repeated 100 times, each with a different set of K participants.

### 4.9 DNN implementation

The DNN was implemented using PyTorch (Paszke et al., 2017) and computed on NVIDIA Titan Xp GPUs using CUDA. More details about hyperparameter tuning are found in Supplementary Methods S1. More details about DNN finetuning are found in Supplementary Methods S2.

### 4.10 Statistical tests

To evaluate whether differences between algorithms were statistically significant, we adapted a bootstrapping approach developed for cross-validation procedures (page 85 of Kuhn and Johnson, 2013). More specifically, we performed bootstrap sampling 1,000 times. For each bootstrap sample, we randomly picked K participants with replacement, while the remaining 10,000 – K participants were used as test participants. Thus, the main difference between our main experiments (100 repeats of K-shot learning in Figure 2A) and the bootstrapping procedure is that the bootstrapping procedure sampled participants with replacement, so the K bootstrapped participants might not be unique. For each of the 1,000 bootstrapped samples, we applied classical (KRR) baseline, basic meta-matching (KRR), basic meta-matching (DNN) and advanced meta-matching (stacking), thus yielding 1,000 bootstrapped samples of COD and Pearson’s correlation (computed from the remaining 10,000 – K participants). Bootstrapping was not performed for advanced meta-matching (finetune) because 1000 bootstrap samples would have required 60 days of compute time (on a single GPU).

Statistical significance for COD and Pearson’s correlation were calculated separately. For ease of explanation, let us focus on COD. The procedure for Pearson’s correlation was exactly the same, except we replaced COD with Pearson’s correlation in the computation. To compute the statistical difference between advanced meta-matching (finetune) and another algorithm X, we first fitted a Gaussian distribution to the 1,000 bootstrapped samples of COD from algorithm X, yielding a cumulative distribution function (CDF_X_). Suppose the average COD of advanced meta-matching (finetune) across the 100 random repeats of K-shot learning was μ. Then the p value was given by 2 * CDF(μ) if μ is less than the mean of the bootstrap distribution, or 2 * (1 - CDF(μ)) if μ is larger than the mean of bootstrap distribution.

When computing the statistical difference between two algorithms X and Y with 1000 bootstrapped samples each, we first fitted a Gaussian distribution to the 1,000 bootstrapped samples of COD from algorithm X, yielding a cumulative distribution function (CDF_X_). This was repeated for algorithm Y, yielding a cumulative distribution function (CDF_Y_). Let the average COD of algorithm X (and Y) across the 100 random repeats of K-shot learning be μ_X_ (and μ_Y_). We can then compute a p value by comparing μ_X_ with CDF_X_ and a p value by comparing μ_Y_ with CDF_Y_. The larger of the two p values was reported.

P values were computed between all pairs of algorithms. Multiple comparisons were corrected using false discovery rate (FDR, q < 0.05). FDR was applied to all K-shots and across all pairs of algorithms

### 4.11 Data and code availability

This study utilized publicly available data from the UK Biobank (https://www.ukbiobank.ac.uk/). Code for the classical (KRR) baseline and meta-matching algorithms can be found here (GITHUB_ LINK). The trained models for meta-matching are available also in the GitHub link above. The code was reviewed by one of the co-authors (LA) before merging into the GitHub repository to reduce the chance of coding errors.

## Acknowledgment

We like to thank Christine Annette, Taimoor Akhtar, Li Zhenhua for their help on the HORD algorithm. This work was supported by the Singapore National Research Foundation (NRF) Fellowship (Class of 2017). Any opinions, findings and conclusions or recommendations expressed in this material are those of the author(s) and do not reflect the views of National Research Foundation, Singapore. Our research also utilized resources provided by the Center for Functional Neuroimaging Technologies, P41EB015896 and instruments supported by 1S10RR023401, 1S10RR019307, and 1S10RR023043 from the Athinoula A. Martinos Center for Biomedical Imaging at the Massachusetts General Hospital. Our computational work was partially performed on resources of the National Supercomputing Centre, Singapore (https://www.nscc.sg). The Titan Xp GPUs used for this research were donated by the NVIDIA Corporation. D.B. was supported by the Healthy Brains Healthy Lives initiative (Canada First Research Excellence fund), the CIFAR Artificial Intelligence Chairs program (Canada Institute for Advanced Research), and Google (Research Award), as well as by NIH grant R01AG068563A. A.J.H. was supported by the National Institutes of Health (Grant R01MH120080). This research has been conducted using the UK Biobank resource under application 25163.

## Supplementary Materials

This supplemental material is divided into *Supplemental Methods* and *Supplemental Results* to complement the Methods and Results sections in the main text, respectively.

## Supplementary Methods

This section provides additional implementation details of the meta-matching. Section S1 provides details about meta-matching with DNN. Section S2 provides details about meta-matching with DNN finetuning.

### S1. Details about basic meta-matching (DNN)

In this section, we provide implementation details of DNN, which we utilized for basic meta-matching (DNN), as well as both advanced meta-matching algorithms.

- The DNN we considered is a generic feedforward neural network, which was implemented with default libraries (class “nn.Linear”) in PyTorch (Paszke et al., 2017).
- The loss function was MSE (mean squared error) loss. The output layer has 33 nodes, which is the number of training meta-set non-brain-imaging phenotypes (phenotypes).
- We used the HORD algorithm (Regis and Shoemaker, 2013; Ilievski et al., 2017; Eriksson et al., 2019) to automatically tune the hyperparameters using the validation set (N = 5370) within the training meta-set. By setting a specific search range for multiple hyperparameters, the HORD algorithm was able to tune these hyperparameters within these ranges automatically. HORD does not perform well when there are too many hyperparameters to tune. Therefore, several hyperparameters were set based on our manual tuning using the training meta-set. These hyperparameters were stochastic gradient descent (SGD) with 0.9 momentum, 128 for batch size and Xavier uniform for weight initialization.
- Table S3 shows the search ranges of hyperparameters tuned by the HORD algorithm. We ran 200 HORD evaluation rounds. For each HORD evaluation round, 1000 epochs were run. DNN was trained on the training set (within the training meta-set) and evaluated on the validation set (within the training meta-set) for each epoch. The epoch with the best COD on the validation set was chosen as the optimal epoch.
- Table S4 shows the final set of hyperparameters estimated by the HORD algorithm. The final DNN structure is a 4-layer DNN. The optimal epoch on the validation set is 118 epochs. After we obtained the best DNN on the training meta-set, we applied the trained DNN to the test meta-set.

**Table S3.** Search ranges of hyperparameters tuned by the HORD algorithm.

| Hyperparameter tuned | Range |
| --- | --- |
| Number of layers | 2 to 5 |
| Number of nodes for each layer (separately) | 2 to 512 |
| Dropout rate | 0 to 0.8 |
| Starting learning rate | 1e-2 to 1e-4 |
| Epochs to decrease the learning rate | 10 to 1000 |
| Weight decay rate | 1e-3 to 1e-7 |

**Table S4.** Final DNN hyperparameters estimated by the HORD algorithm.

| Hyperparameter | Value |
| --- | --- |
| Number of layers | 4 |
| Number of nodes for each layer (separately) | 87/386/313/33 |
| Dropout rate | 0.242 |
| Starting learning rate | 3.646e-03 |
| Epochs to decrease learning rate | 312 |
| Weight decay rate | 8.447e-04 |

### S2. Details about advanced meta-matching (finetune)

In this section, we provide implementation details of advanced meta-matching (finetune). The trained DNN (previous section) was applied to the K participants in the test meta-set. For a given test meta-set phenotype,

- The best DNN output that gave the best prediction for the test phenotype (based on the K participants) was selected
- We took the trained DNN and removed all output nodes except the best DNN output node (selected in the previous step). We then performed finetuning on this DNN using the K participants. The loss function was MSE (mean squared error) loss. The evaluation metric was COD.
- Finetuning was only performed on the weights of the last two layers. The weights of the earlier layers were frozen. We split the K subjects into training and validation sets (4:1 ratio). We ran the finetuning for 100 epochs using the training set and checked the performance in the validation set every 10 epochs. The DNN from the epoch with the best performance in the validation set was used for predicting the phenotype in the remaining 10,000 – K participants. If the performance in the validation set was worse than the original DNN (without finetuning), then we simply applied the original DNN to the remaining 10,000 – K participants. We did not perform cross-validation like the classical (KRR) baseline, because the runtime would be increased multiple folds.
- Furthermore, because of the small number of participants K, we decided not to optimize the hyperparameters of the finetuning procedure for fear of overfitting. Optimizing the hyperparameters would also be computationally too expensive. More specifically, it took 6 days (on one GPU) to run 34 meta-set phenotypes for 100 repetition of K-shots across different values of K. Optimizing the hyperparameters using HORD would dramatically increase the runtime to 6 x 200 = 1200 days (since we utilized 200 HORD rounds).
- Therefore, we simply set the hyperparameters to the following generic values: stochastic gradient descent (SGD) with 0.9 momentum. The learning rate was set to be 1e-3. The batch size was set to be the minimum of K and 32. So if K was less than 32, the batch size was set to be K. Otherwise, the batch size was set to be 32.

## Supplementary Results

**Table S1.** Dictionary of 33 training meta-set non-brain-imaging phenotypes. For UK Biobank IDs, please see GITHUB_LINK.

| Label | Description |
| --- | --- |
| ECG C1 | ECG measures principal component 1 |
| Sex | sex |
| Sex G C2 | genotype sex inference principal component 2 |
| Body C2 | anthropometry principal component 2 |
| Grip C1 | hand grip strength principal component 1 |
| Body C1 | anthropometry principal component 1 |
| Bone C3 | bone-densitometry of heel principal component 3 |
| BP eye C4 | blood pressure & eye measures principal component 4 |
| Matrix C1 | matrix pattern completion principal component 1 |
| #Mem C1 | numeric memory principal component 1 |
| Matrix C2 | matrix pattern completion principal component 2 |
| Fluid Int. | fluid intelligence |
| Hearing | hearing signal-to-noise-ratio (snr) of triplet (left) |
| Illness C1 | non-cancer illness principal component 1 |
| #household | number of people in household |
| Time TV | time spent watching television (tv) per day |
| BP eye C2 | blood pressure & eye measures component 2 |
| Body C3 | anthropometry principal component 3 |
| ECG C6 | ECG measures principal component 6 |
| ECG C2 | ECG measures principal component 2 |
| Illness C4 | non-cancer illness principal component 4 |
| Smoke C1 | smoke principal component 1 |
| BP eye C3 | blood pressure & eye measures principal component 3 |
| BP eye C6 | blood pressure & eye measures principal component 6 |
| Urine C1 | urine assays principal component 1 |
| Sex G C1 | genotype sex inference principal component 1 |
| Bone C1 | bone-densitometry of heel principal component 1 |
| Matrix C3 | matrix pattern completion principal component 3 |
| Time walk | number of days walked 10+ minutes per week |
| BP eye C5 | blood pressure & eye measures principal component 5 |
| ECG C3 | ecg measures principal component 3 |
| Genetic C1 | genetic principal components and heterozygosity principal component 1 |
| Sleep | sleep duration per day |

**Table S2.** Dictionary of 34 test meta-set non-brain-imaging phenotypes. For UK Biobank IDs, please see GITHUB_LINK.

| Label | Description |
| --- | --- |
| Alcohol 3 | average weekly beer plus cider intake |
| Blood C2 | blood assays principal component 2 |
| Breath C1 | spirometry principal component 1 |
| Age | age |
| Cancer C1 | cancer principal component 1 |
| Carotid C1 | carotid ultrasound principal component 1 |
| Match-o | pairs matching online |
| Trail C1 | trail making principal component 1 |
| Digit-o C1 | symbol digit substitution online principal component 1 |
| Digit 1 | symbol digit substitution principal component 1 |
| Match | pairs matching |
| ProMem C1 | prospective memory principal component 1 |
| RT C1 | reaction time principal component 1 |
| Trail-o C1 | trail making online principal component 1 |
| Tower C1 | tower rearranging principal component 1 |
| Family C1 | family history (parent's age) principal component 1 |
| Blood C5 | blood assays principal component 5 |
| Dur C4 | process durations principal component 4 |
| Dur C2 | process durations principal component 2 |
| Loc C1 | location principal component 1 |
| Dur C1 | process durations principal component 1 |
| Digit-o C6 | symbol digit substitution online principal component 6 |
| Trail-o C4 | trail making online principal component 4 |
| Blood C4 | blood assays principal component 4 |
| Alcohol 2 | average weekly champagne plus white wine intake |
| Carotid C5 | carotid ultrasound principal component 5 |
| Time drive | time spent driving per day |
| Travel | frequency of travelling from home to job workplace per week |
| Work | weekly length of working hour for main job |
| Age edu | age completed full time education |
| Deprive C1 | multiple deprivation principal component 1 |
| Blood C3 | blood assays principal component 3 |
| Alcohol 1 | average monthly spirits intake |
| Neuro | neuroticism score |

**Figure S1.**
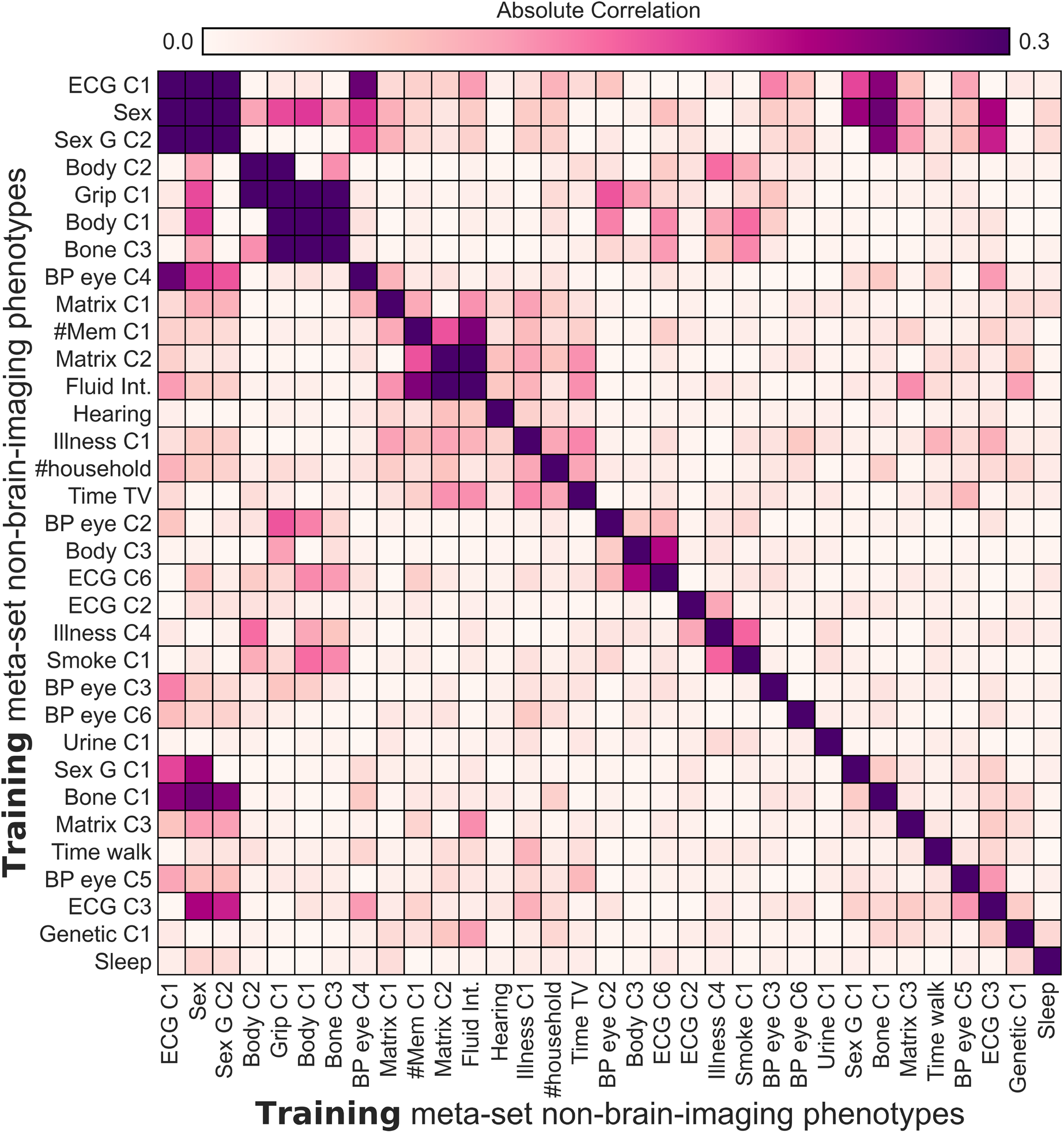
Absolute Pearson’s correlation among 33 non-brain-imaging phenotypes in the training meta-set.

**Figure S2.**
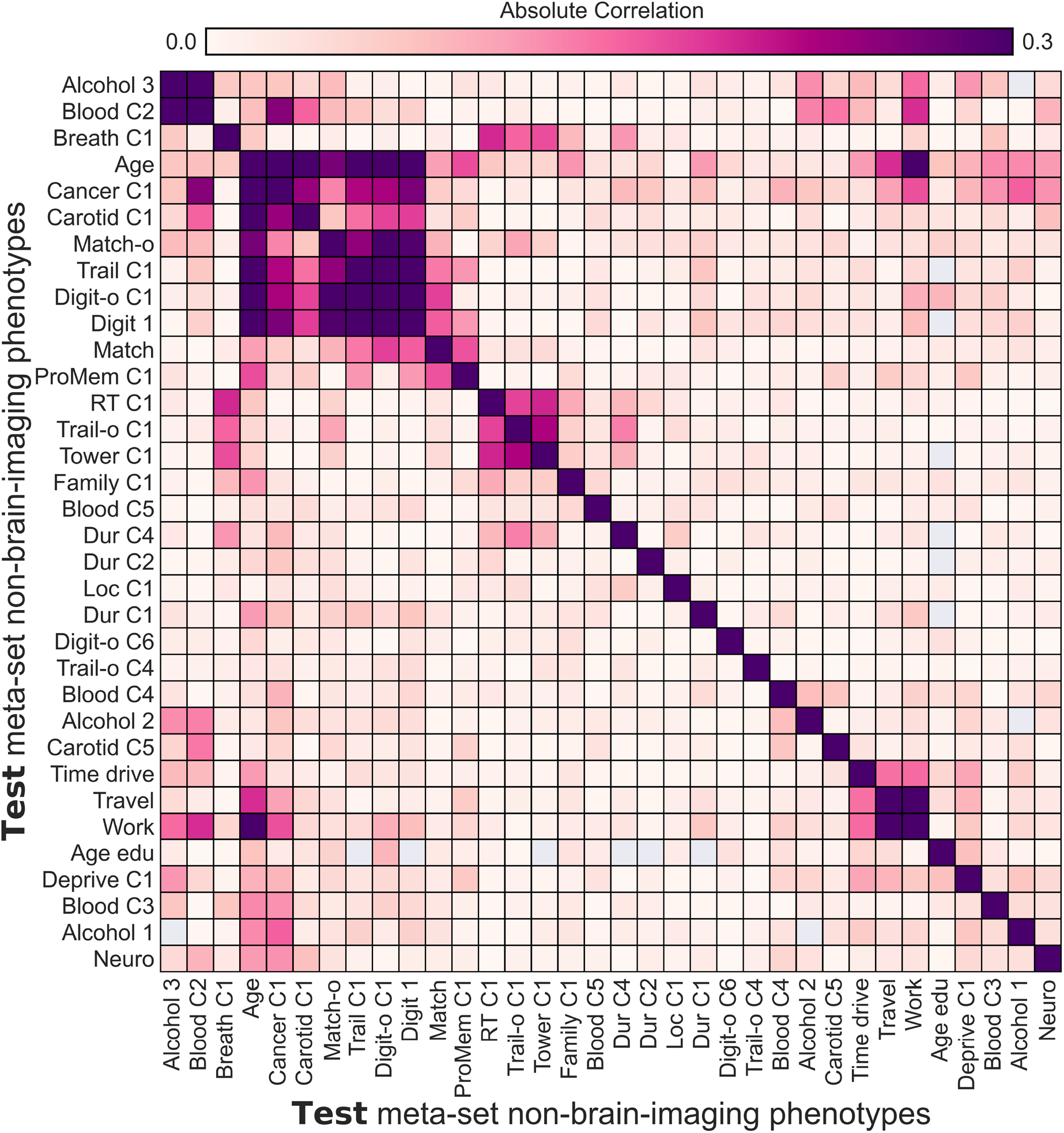
Absolute Pearson’s correlation among 34 non-brain-imaging phenotypes in the test meta-set.

**Figure S3.**
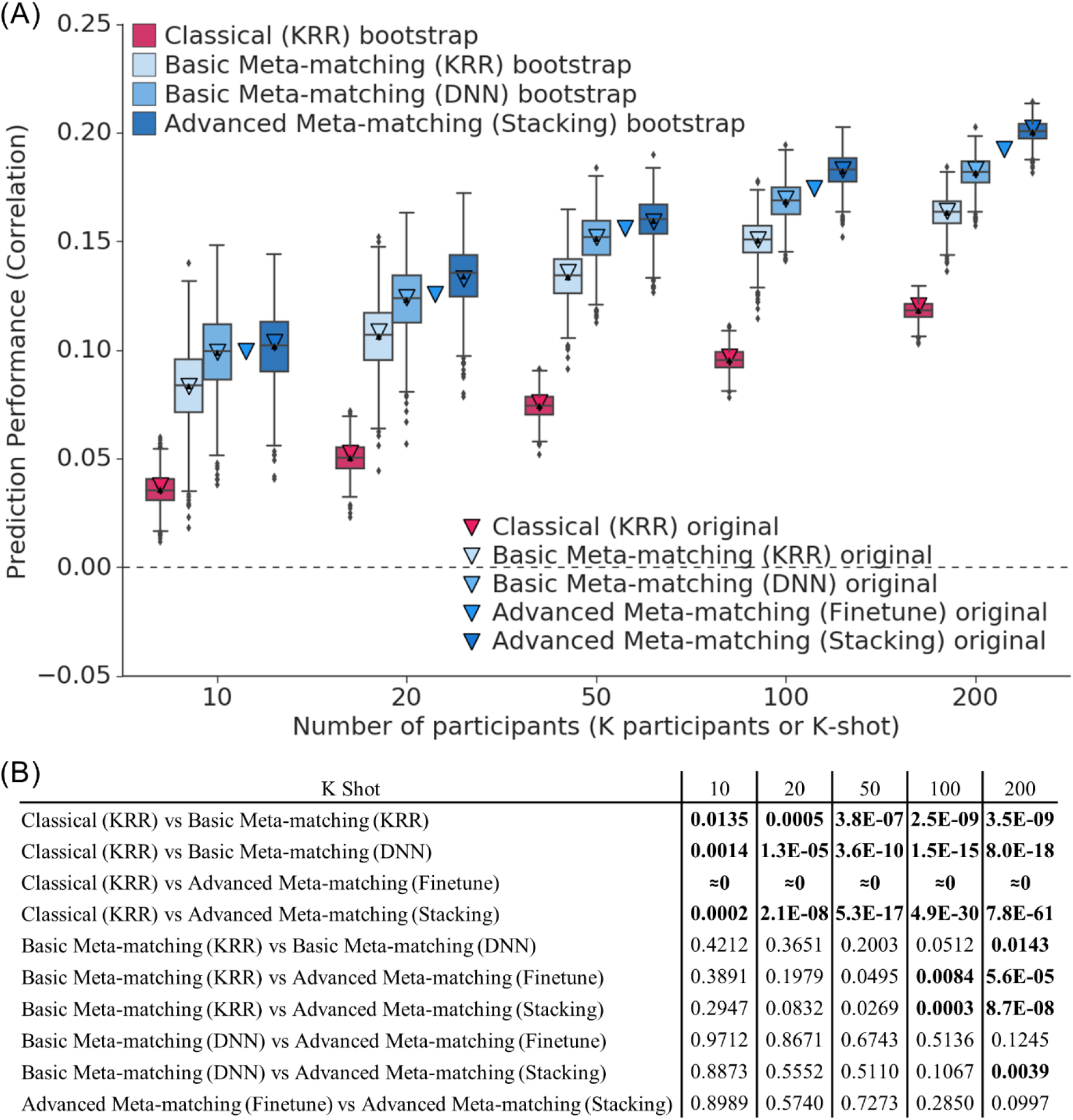
Meta-matching outperformed classical kernel ridge regression (KRR) baseline. (A) Prediction performance (Pearson’s correlation) with different number of participants. This plot is the same as Figure 4A, but the boxplots now show the bootstrap distribution of each approach based on 1000 bootstrapped samples. The triangles show the average performance (Pearson’s correlation) of 34 non-brain-imaging phenotypes using the original 100 random repeats (Figure 4A). We observe that the mean of the bootstrap distributions matches the mean of the original experiments (Figure 4A) quite well. Bootstrapping could not be performed for advanced meta-matching (finetune) because 1000 bootstrap samples would have required 60 days of compute time. (B) Statistical differences among the different algorithms. For rows comparing advanced meta-matching (finetune) and another algorithm X, p values were derived by comparing the mean of advanced meta-matching (finetune) with algorithm X’s bootstrap distribution (assuming Gaussanity). For other rows comparing algorithms X and Y, bootstrap distributions were available for both X and Y. Therefore, one p value was obtained by comparing the original mean of X with Y’s bootstrap distribution and another p value was obtained by comparing the original mean of Ywith X’s bootstrap distribution. The larger of the two p values were reported. Bold indicates statistical significance after FDR correction (q < 0.05).

**Figure S4.**
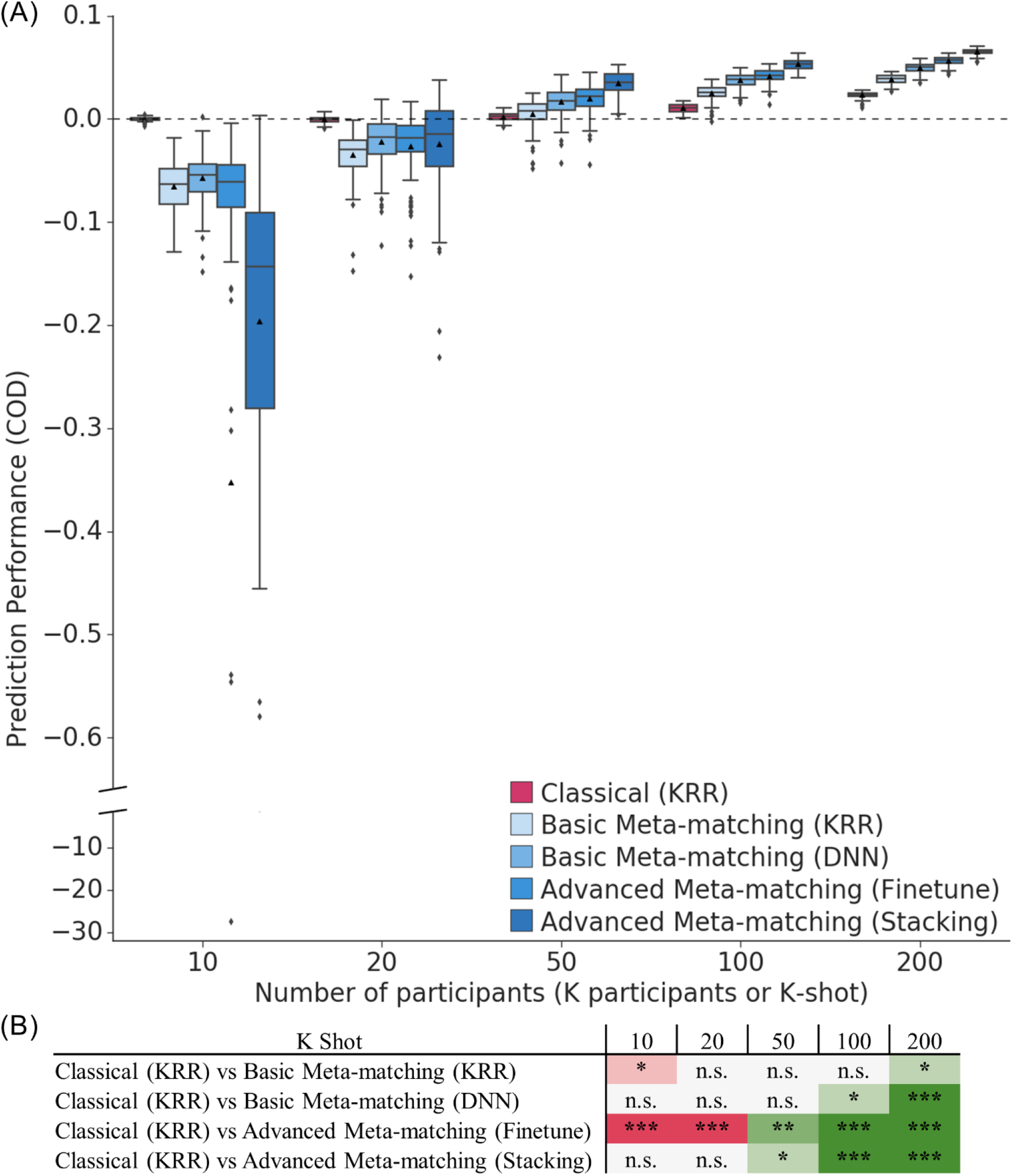
Meta-matching outperformed classical kernel ridge regression (KRR) baseline. (A) Prediction performance (coefficient of determination; COD) averaged across 34 non-brain-imaging phenotypes in the test meta-set (N = 10,000 – K). The K participants were used to train and tune the models (Figure 3). Boxplots represent variability across 100 random repeats of K participants (Figure 2A). Whiskers represent 1.5 inter-quartile range. (B) Statistical difference between the prediction performance (COD) of classical (KRR) baseline and meta-matching algorithms. “n.s.” indicates that difference was not statistically significant after multiple comparisons correction (FDR q < 0.05). “*” indicates p < 0.05 and statistical significance after multiple comparisons correction (FDR q < 0.05). “**” indicates p < 0.001 and statistical significance after multiple comparisons correction (FDR q < 0.05). “***” indicates p < 0.00001 and statistical significance after multiple comparisons correction (FDR q < 0.05). Green indicates that meta-matching outperforms classical (KRR) baseline. Red indicates that classical (KRR) baseline outperforms meta-matching. Observe that all algorithms performed poorly (COD ≤ 0) when there were less than 50 participants (K < 50), suggesting chance or worse than chance prediction for all algorithms.

**Figure S5.**
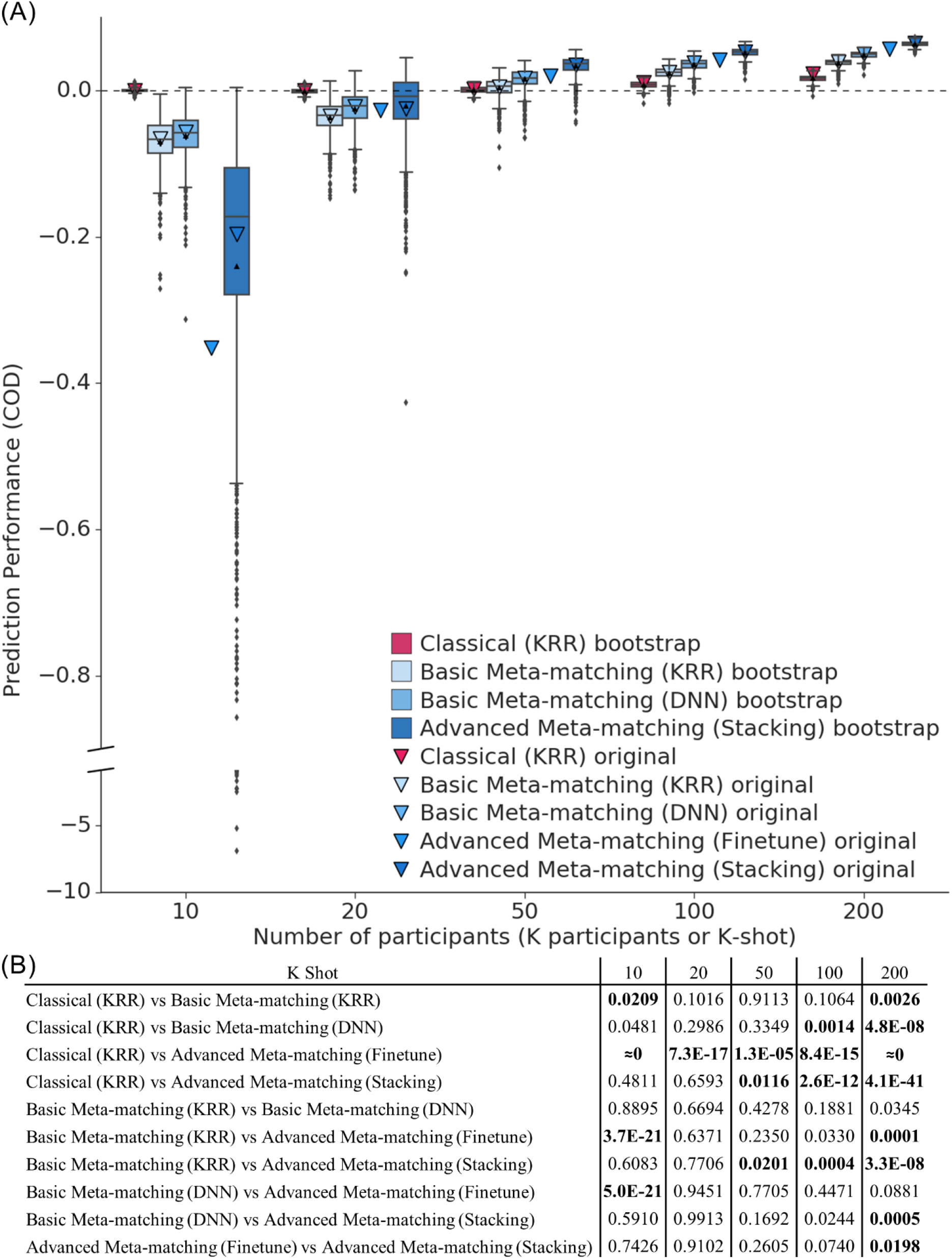
Meta-matching outperformed classical kernel ridge regression (KRR) baseline. (A) Prediction performance (coefficient of determination; COD) with different number of participants. This plot is the same as Figure S4A, but the boxplots now show the bootstrap distribution of each approach based on 1000 bootstrapped samples. The triangles show the average performance (COD) of 34 non-brain-imaging phenotypes using the original 100 random repeats (Figure S4A). We observe that the mean of the bootstrap distributions matches the mean of the original experiments (Figure S4A) quite well. Bootstrapping could not be performed for advanced meta-matching (finetune) because 1000 bootstrap samples would have required 60 days of compute time. (B) Statistical differences among the different algorithms. For rows comparing advanced meta-matching (finetune) and another algorithm X, p values were derived by comparing the mean of advanced meta-matching (finetune) with algorithm X’s bootstrap distribution (assuming Gaussanity). For other rows comparing algorithms X and Y, bootstrap distributions were available for both X and Y. Therefore, one p value was obtained by comparing the original mean of X with Y’s bootstrap distribution and another p value was obtained by comparing the original mean of Y with X’s bootstrap distribution. The larger of the two p values were reported. Bold indicates statistical significance after FDR correction (q < 0.05).

**Figure S6.**
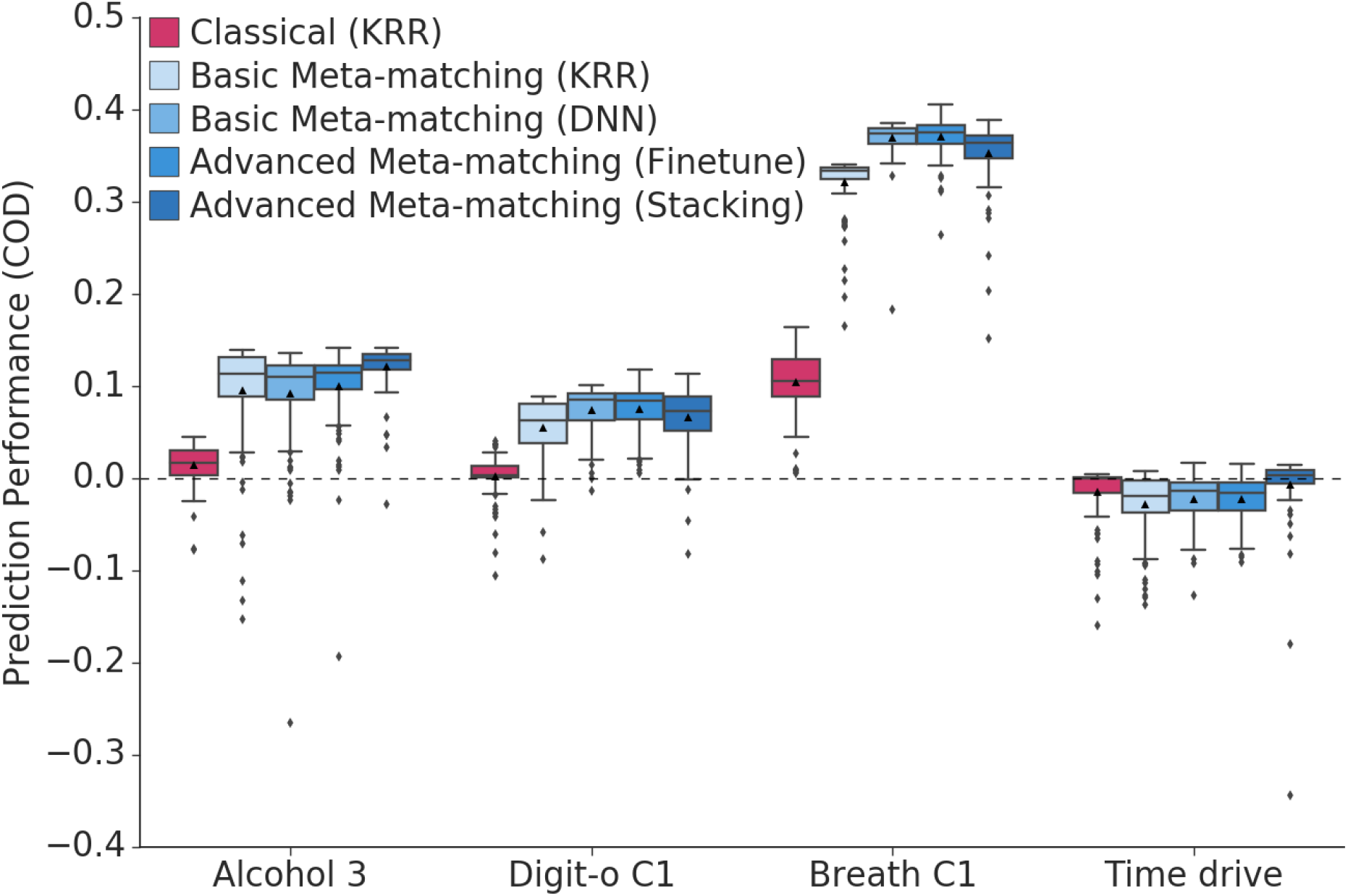
Examples of non-brain-imaging phenotypic prediction performance in the test meta-set in the case of 100-shot learning. Here, prediction performance was measured using coefficient of determination (COD). “Alcohol 3” (average weekly beer plus cider intake) was most frequently matched to “Bone C3” (bone-densitometry of heel principal component 3). “Digit-o C1” (symbol digit substitution online principal component 1) was most frequently matched to “Matrix C1” (matrix pattern completion principal component 1). “Breath C1” (spirometry principal component 1) was most frequently matched to “Grip C1” (hand grip strength principal component 1). “Time drive” (Time spent driving per day) was most frequently matched to “BP eye C3” (blood pressure & eye measures principal component 3).

**Figure S7.**
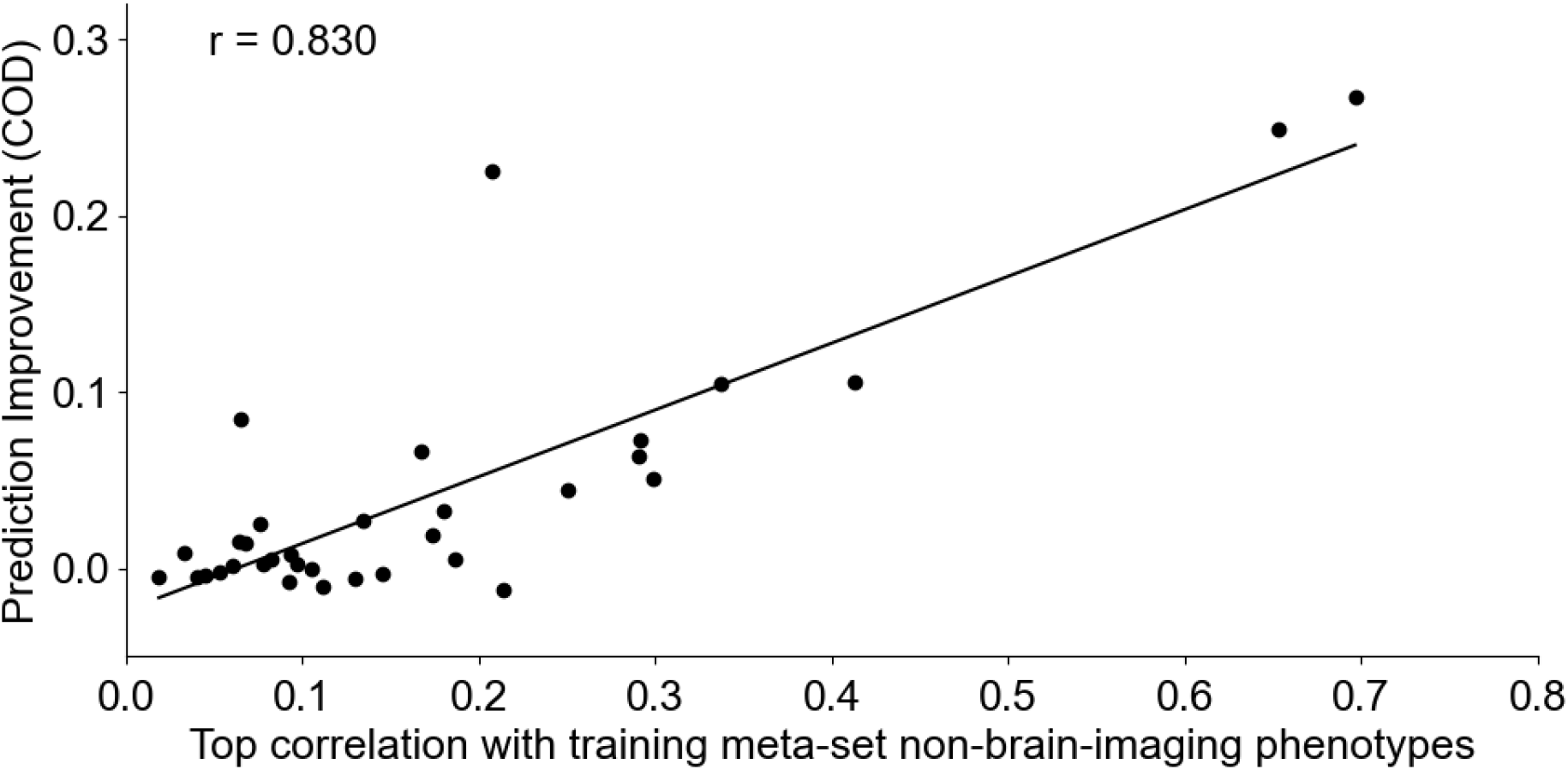
Prediction improvements were driven by correlations between training and test meta-set phenotypes. Vertical axis shows the prediction improvement of advanced meta-matching (stacking) with respect to classical (KRR) baseline under the 100-shot scenario. Prediction performance was measured using coefficient of determination (COD). Each dot represents a test meta-set phenotype. Horizontal axis shows each test phenotype’s top absolute Pearson’s correlation with phentoypes in the training meta-set. Test phenotypes with stronger correlations with at least one training phenotype led to greater prediction improvement with meta-matching.

**Figure S8.**
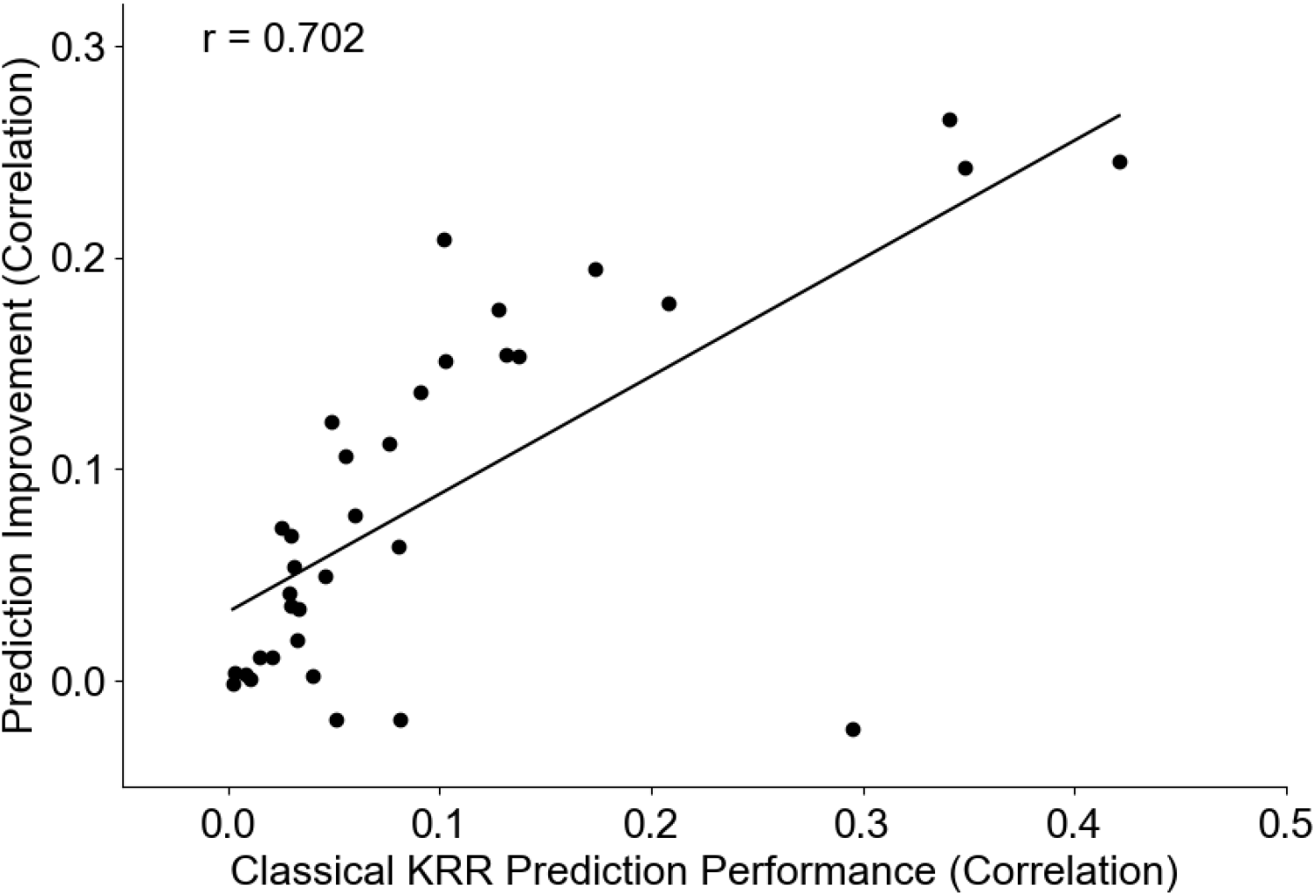
Phenotypes better predicted by classical kernel ridge regression benefited more from meta-matching. Vertical axis shows the prediction improvement of advanced meta-matching (stacking) with respect to classical (KRR) baseline under the 100-shot scenario. Prediction performance was measured using Pearson’s correlation. Each dot represents a test meta-set phenotype. Horizontal axis shows the prediction performance with the classical (KRR) baseline under the 100-shot scenario. Similar conclusions were obtained with coefficient of determination (Figure S9).

**Figure S9.**
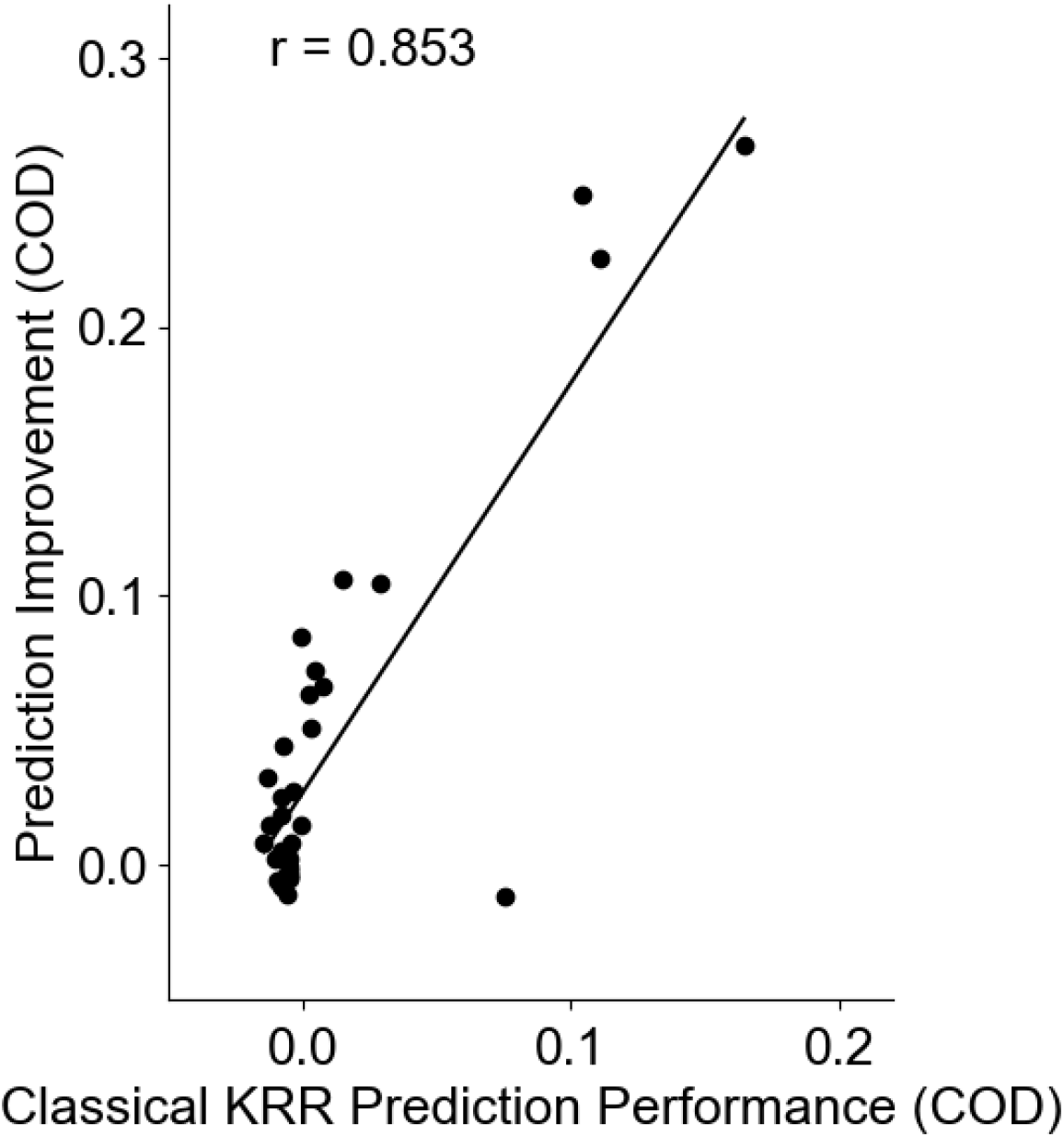
Phenotypes better predicted by classical kernel ridge regression benefited more from meta-matching. Vertical axis shows the prediction improvement of advanced meta-matching (stacking) with respect to classical (KRR) baseline under the 100-shot scenario. Prediction performance was measured using coefficient of determination (COD). Each dot represents a test meta-set phenotype. Horizontal axis shows the prediction performance with the classical (KRR) baseline under the 100-shot scenario.

